# RAB18 confers protection to lipid droplets from lysolipophagy and counters cellular steatosis

**DOI:** 10.1101/2023.03.11.531838

**Authors:** Adrian Rieck, Georgia Guenther, Jan G. Hengstler, Nachiket Vartak

## Abstract

Lipid droplet (LD) enlargement is the most conspicuous cellular phenotype of steatotic liver diseases such as NAFLD and NASH. The small GTPase RAB18 localizes to lipid droplets yet its effects on LD dynamics are unknown. We show that RAB18 localizes to LDs dynamically via the acylation cycle in an activity dependent manner. Removal of RAB18 from LDs through acylation cycle inhibition of RNAi-mediated downregulation leads to an increase in LD size, that can be prevented by inhibition of autophagy. We propose a model where RAB18 on LDs decreases the susceptibility of LDs to autophagy by lysosomes, allowing them to exist longer in cells and enabling an even distribution of exogenous lipid load. In the absence of RAB18, fewer LDs survive autophagic removal, and enlarge disproportionately as they absorb the incumbent exogenous lipid load on cells – leading to cellular steatosis. Inhibition of autophagy prevented cellular steatosis in primary human hepatocytes. Finally, mice treated with the autophagy inhibitor chloroquine were resistant to the development of liver steatosis when on a steatogenic diet. The modulation of RAB18 activity and autophagy inhibition represents a promising approach to prevent steatotic liver disease in the clinical setting.

**Graphical Abstract:** 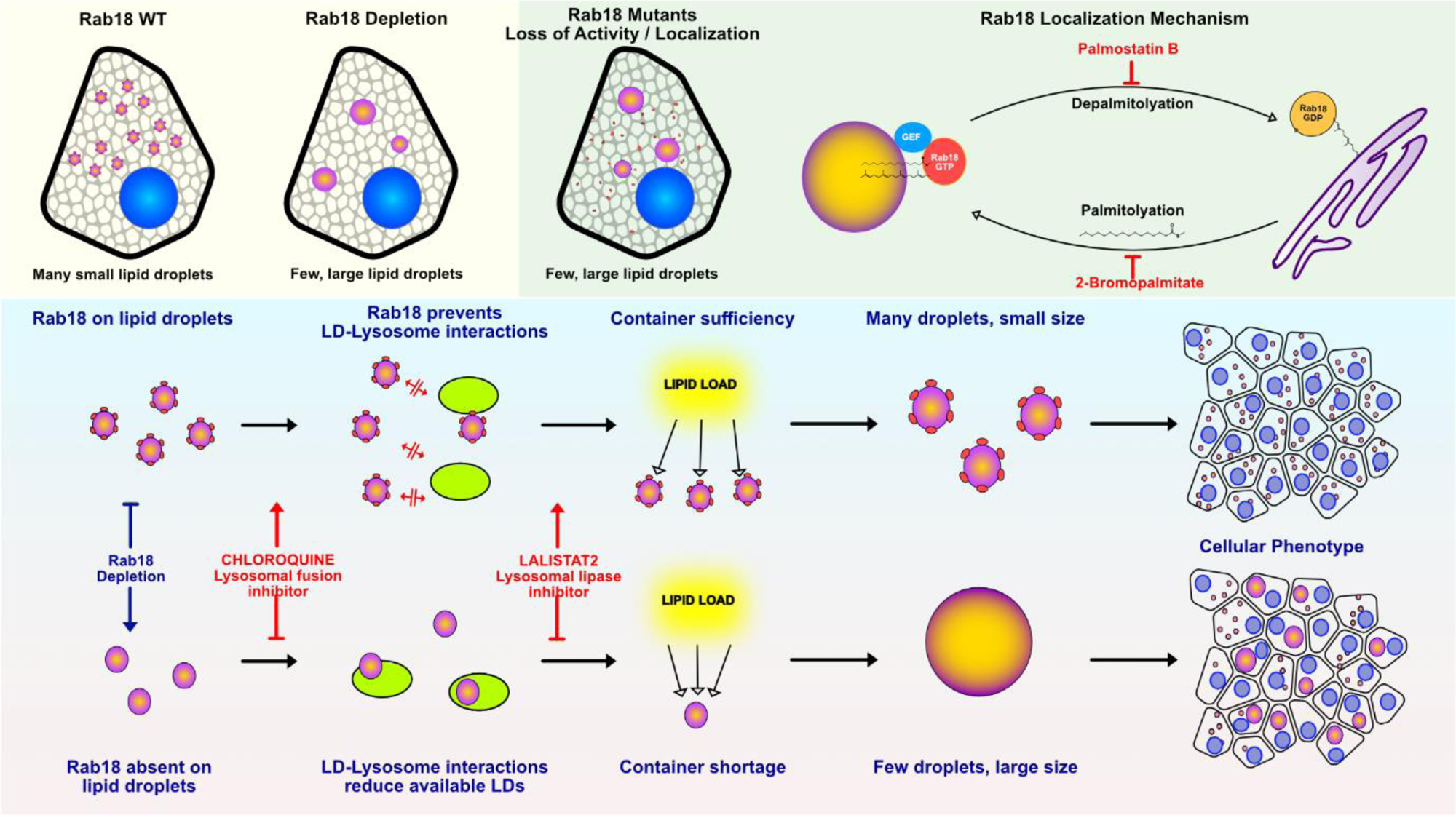

**One line summary:** RAB18 prevents lipid droplet autophagy to distribute lipid load in cells and prevent steatosis.

## Introduction

Lipid droplets (LDs) are single-membraned organelles derived from the endoplasmic reticulum, which are used as storage reservoirs for neutral lipids. Adipocytes, hepatocytes, and intestinal epithelial cells convert free fatty acids to triacylglycerol (TAGs) and import them into lipid droplets for safe storage. In adipocytes, lipid droplets may reach sizes comparable to the cell itself as part of normal physiology^1^. In the liver, however, abnormally large lipid droplets in hepatocytes represent pathological steatosis as seen in metabolic-associated fatty liver disease (MAFLD)^2^. Beginning as dysregulation of lipid storage in LDs, MAFLD eventually progresses to steatohepatitis, cirrhosis, and hepatocellular carcinoma. Despite their conspicuous role in metabolic liver disease, the molecular processes that regulate lipid droplet size in hepatocytes remain unknown.

Lipid droplet size is regulated through the activity of several LD-and ER-resident proteins such as PLIN2, NRZ and DGAT^3–5^. However, the Rab family of small GTPases is the canonical regulator of membrane bound organelles^6^ and it is the small GTPase RAB18 that imparts the lipid droplet its identity. In adipocytes, it is known that depletion of RAB18 leads to lipid droplet enlargement^7,8^. It is however not known how this effect is mediated in adipocytes, nor if it has any role in lipid droplet size regulation in hepatocytes.

Proteomic profiling of lipid droplets in hepatocytes under exogenous lipid load shows no changes in RAB18 expression under conditions where LDs enlarge^9,10^. This is not surprising, as the function of small GTPases is regulated mainly through the nucleotide-loading and their localization to appropriate target membranes. Like all small GTPases^6^, RAB18 switches between GTP-bound, effector-binding ‘active’ state and a GDP-bound ‘inactive’ state. RAB18[GTP] converts to RAB18[GDP] through interaction with GTPase-activating proteins (GAPs), while the GDP can be re-exchanged for GTP through interaction with guanine-nucleotide exchange factors (GEFs). Interconversion between these two states of RAB18 through the ‘GTPase’ cycle allows RAB18 to function as molecular switch.

Besides the common GTPase cycle, the localization of small GTPases to target membranes is subject to an additional layer of regulation through post-translational prenylation^11^ and palmitoylation^12^. The C-termini of GTPases contain a specific amino acid sequence called a CAAX box^13^, where the Cys residue is post-translationally S-prenylated. Hydrophobic irreversible prenylation at the C-terminus confers non-specific membrane affinity. Guanine dissociation inhibitors (GDIs) can bind and shield the hydrophobic prenyl group to solubilize these membrane-bound Rab proteins to the cytosol based on their GTP-loading state. Finally, the membrane affinity of prenylated Rab proteins can be modulated by additional palmitoylation of cysteines in their C-terminus. Compartment-specific palmitoylation adds hydrophobicity to Rab proteins, thus increasing their membrane affinity and residence time on target membranes. Ubiquitous cell-wide depalmitoylation occurs via acyl-protein thioesterases. Cell-wide depalmitoylation and target-membrane-specific palmitoylation results in an ‘acylation cycle’ wherein Rab proteins are enriched but dynamically turned over on their target membrane. These reaction-diffusion cycles, rather than protein expression, typically regulate the activity of small GTPases.

Differences exist in the GTPase and acylation cycles of the Rab family, depending on the inherent GTPase catalytic activity, and presence of appropriate cysteine residues at the C-terminus. Whether RAB18 undergoes an acylation cycle, and how it influences lipid droplet size remains as yet undiscovered.

In this work, we leverage live cell confocal fluorescence and coherent anti-stokes Raman (CARS) microscopy to show that RAB18 localization to the LD membrane is dynamic and modulated by the acylation cycle. Using mutagenic and pharmaceutical perturbations of RAB18 activity, we discover that RAB18 prevents lipid droplet enlargement in hepatocytes by preventing their degradation by lysosomes. We develop the ‘container-shortage’ concept that explains this seemingly counter-intuitive finding – wherein reducing degradation paradoxically also reduces enlargement. Application of this concept in primary human hepatocytes and a mouse model of liver steatosis led to a translationally relevant strategy to constrain pathological lipid droplet enlargement as seen in liver diseases such as MAFLD.

## Materials and Methods

Materials and methods for plasmids and molecular biology, cell culture, Western blotting, confocal fluorescence and CARS imaging and mouse studies is provided in the Supporting Information.

## Results

### RAB18-depletion leads to LD enlargement, which is rescued in an activity and LD-localization dependent manner

Depletion of RAB18 through RNAi in HepG2 cells under exogenous lipid load leads to an enlargement of lipid droplets over period of 20h (**Fig 1A, Supplementary movie M1**), confirming that RAB18 regulates droplet size in liver cells, like in adipocytes. Timelapse imaging of lipid droplet growth shows that is a continuous process which generates a broad distribution of lipid droplet sizes, with only a relatively small number reaching extremely large sizes. The effect of interventions such as RAB18 depletion is reflected more strongly in the upper percentiles of lipid droplet size, while the lower percentiles show a relatively small effect. This is illustrated in the associated histograms, where the RAB18-depletion leads to a 1.5-fold increase at the 75th and nearly a 2-fold increase 99th percentile of lipid droplets diameter, while the mean and median size increases only 1.2-fold. We therefore developed image processing pipelines based on manual segmentation, as well as machine-learning based segmentation (StarDist^14^) to measure lipid droplet size accurately in further experiments.

**Figure 1.**
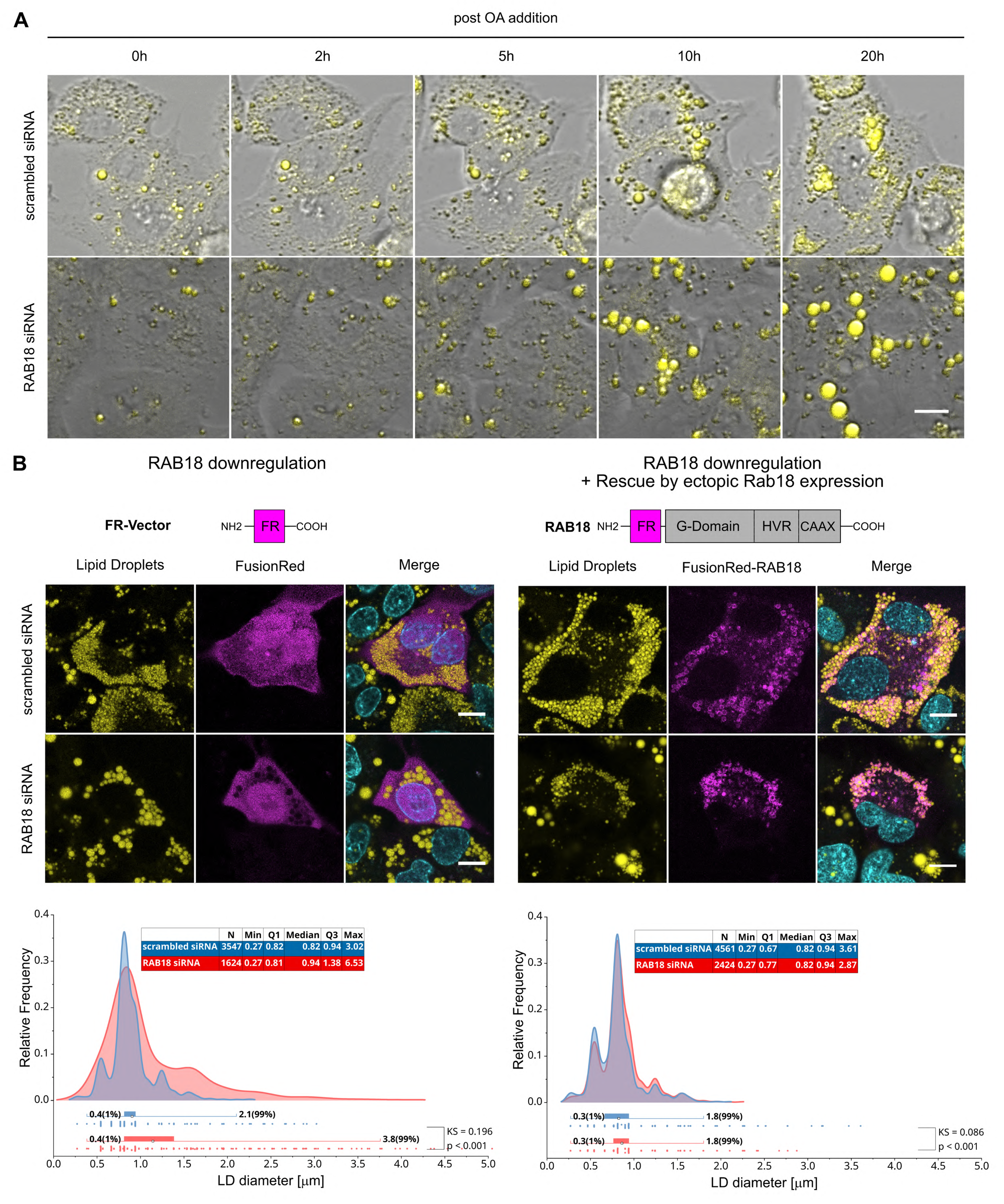
RAB18 localization is essential to rescue WT after RAB18 depletion. **A.** Representative time series showing LD formation stained with BODIPY 493/503 over 20h of OA exposure. RAB18 downregulation leads to an increase in LD size **B.** Representative images of HepG2 cells expressing FusionRed constructs along with LDs visualized with BODIPY. Ectopic expression of FusionRed-RAB18 (right panel), but not FusionRed alone (left panel) rescues the LD-size increase caused by RAB18 depletion after OA-induction. Graph shows the kernel density estimate of the distribution of LDs, while underlying bars indicate quantiles and statistical moments. Number of experiments (N) and Total number of LDs quantified (n): (n, N _FR-scrambled_ = 3547, 4; n, N_FR-siRNA_ = 1624,4; n, N_RAB18-scrambled_=4561, 3; n, N_RAB18-siRNA_=2424, 3. Contrast maximized for display. Scalebar: 10 µm

Ectopic transient expression of fluorescently tagged RAB18 showed that FusionRed-RAB18, but not FusionRed alone, localized to lipid droplets (**Fig 1B**). In cells expressing FusionRed-RAB18, LD-enlargement due endogenous RAB18-depletion was rescued. Two canonical activity modifying mutants are known for small GTPases such as RAB18: Q67L which is constitutively active^15^, and S22N which is constitutively inactive. FusionRed-RAB18Q67L localized to lipid droplets similar to wild-type RAB18 and rescued the LD-enlargement due to endogenous RAB18 depletion (**Fig 2A**). In contrast, the inactive FusionRed-RAB18S22N neither localized to lipid droplets, nor rescued LD-enlargement due to endogenous RAB18-depletion **(Fig. 2B)**.

**Figure 2.**
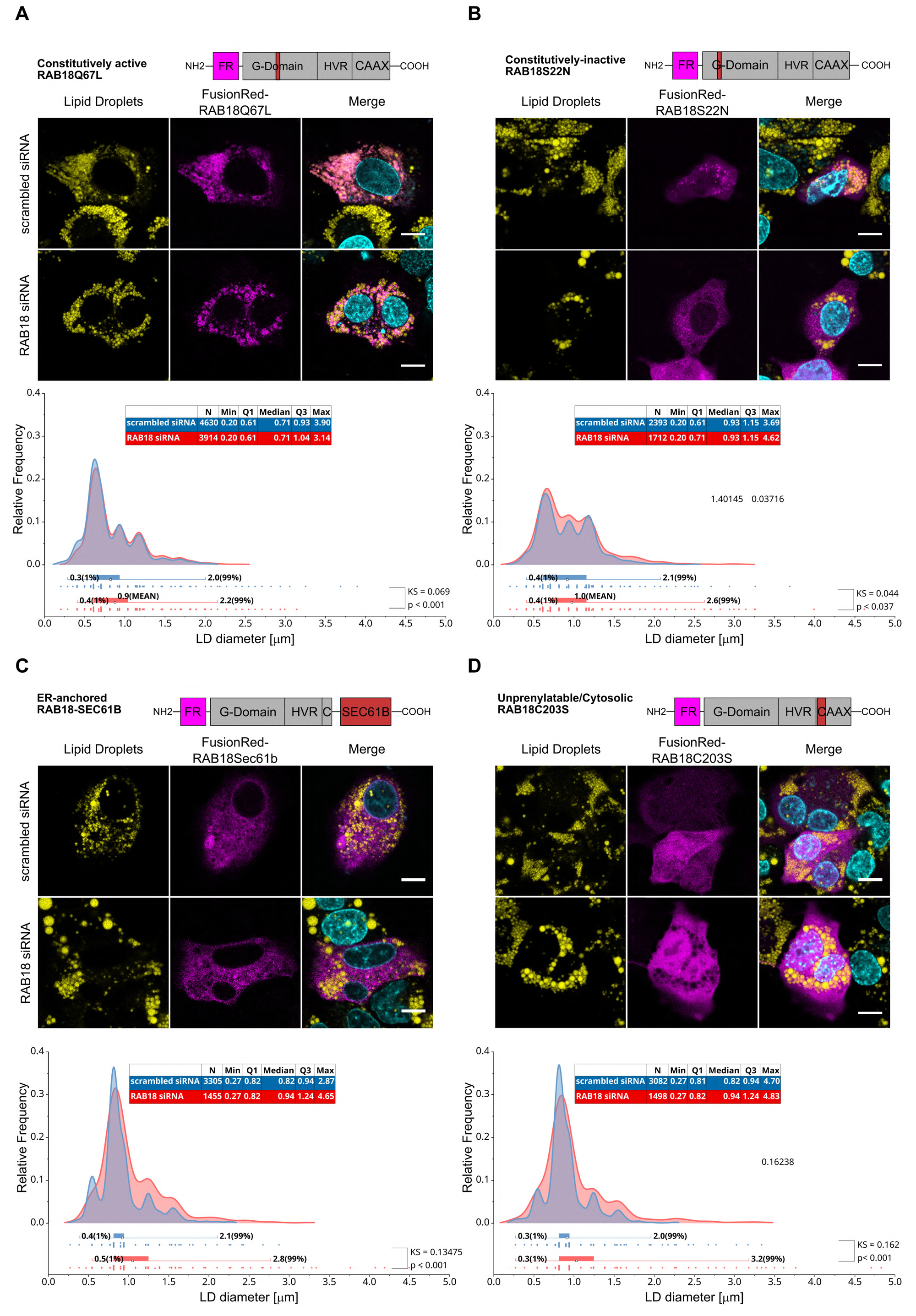
RAB18 activity and localization is essential to rescue WT after RAB18 depletion. **A.** Representative images of HepG2 cells expressing constitutively active (left panel) and dominant negative (right panel) RAB18 mutants as well as the KDE of the LD size distribution. Expression of constitutively active RAB18 rescued the LD size increase caused by RAB18 depletion, while LD size remains increased in cells overexpressing dominant negative RAB18 with endogenous RAB18 depletion. **B.** Representative images of HepG2 cells expressing ER-localized RAB18-Sec61b (left panel) and mutant RAB18 without a prenylation site at C203 (right panel) as well as the KDE of the LD size distribution. Both overexpressed constructs were unable to localize to the LD membrane and failed to rescue the increase in LD size caused byRAB18 depletion. Number of experiments (N) and Total number of LDs quantified (n): n, N_Q67L-scrambled_= 4630, 3; n,N_Q67L-siRNA_= 3914, 3; n,N_S22N-scrambled_= 2393, 3; n,N_S22N-siRNA_= 1712, 3; n,N_Sec61b-scrambled_= 3305, 3; n,N_Sec61b-siRNA_= 1455, 3; n,N_C203S-scrambled_= 3082, 4; n,N_C203S-siRNA_= 1498, 4; Contrast maximized for display. Scalebar: 10 µm

RAB18 is known to localize partly to the ER and participate in lipid droplet-ER contacts during lipid droplet biogenesis^7^. We therefore investigated if the rescue effect under RAB18-depletion is mediated by RAB18 on the ER. However, an ER-targeted FusionRed-RAB18-Sec61b variant, which localized to the ER, could not rescue LD-enlargement due to endogenous RAB18 depletion **(Fig. 2C)**.

Further, RAB18 may be prenylated at the C-terminus, and therefore can exist in a GDI-bound cytosolic state. To investigate if the rescue effect is mediated by an interaction of RAB18 with another cytosolic factor, we ectopically expressed the unprenylatable FusionRed-RAB18-C203S mutant in cells under endogenous RAB18 depletion. This mutant localized to the cytosol as expected, but again, could not rescue LD-enlargement (**Fig 2D**).

Together, these results show that RAB18 must be active to localize to lipid droplets, and only RAB18 that is localized to lipid droplets can prevent lipid droplet enlargement, suggesting a causal link between RAB18 activity, localization and constrained lipid droplet size.

### RAB18 localizes to lipid droplets via C-terminal palmitoylation

The mechanism by which RAB18 localizes to lipid droplets is relevant for lipid droplet size regulation. Small GTPases localize to their target membranes partly through reversible post-translational S-palmitoylation of cysteines in their C-terminal hypervariable regions (HVR). Sequence comparisons of RAB18 with other known S-palmitoylated GTPases^16^ show that the RAB18 HVR contains a potentially palmitoylatable Cys at position 199. Total palmitoylated protein was extracted from HepG2 cells by incubating them with an azide-palmitate analog to be incorporated into palmitoylated proteins. Proteins which had incorporated azide were linked by copper-free Click-conjugation to DBCO-biotin and pulled down with Streptavidin coated beads. Western blot analysis of the pulldown showed that RAB18 is reliably detected in the palmitoylated protein fraction along with other known palmitoylated proteins e.g., RAS (**Fig 3A**), while palmitoylated proteins (e.g., exogenously expressed GFP are absent). The graphs show that RAB18 is enriched approximately 10-fold in the palmitoylated proteins fraction compared to total protein.

**Figure 3.**
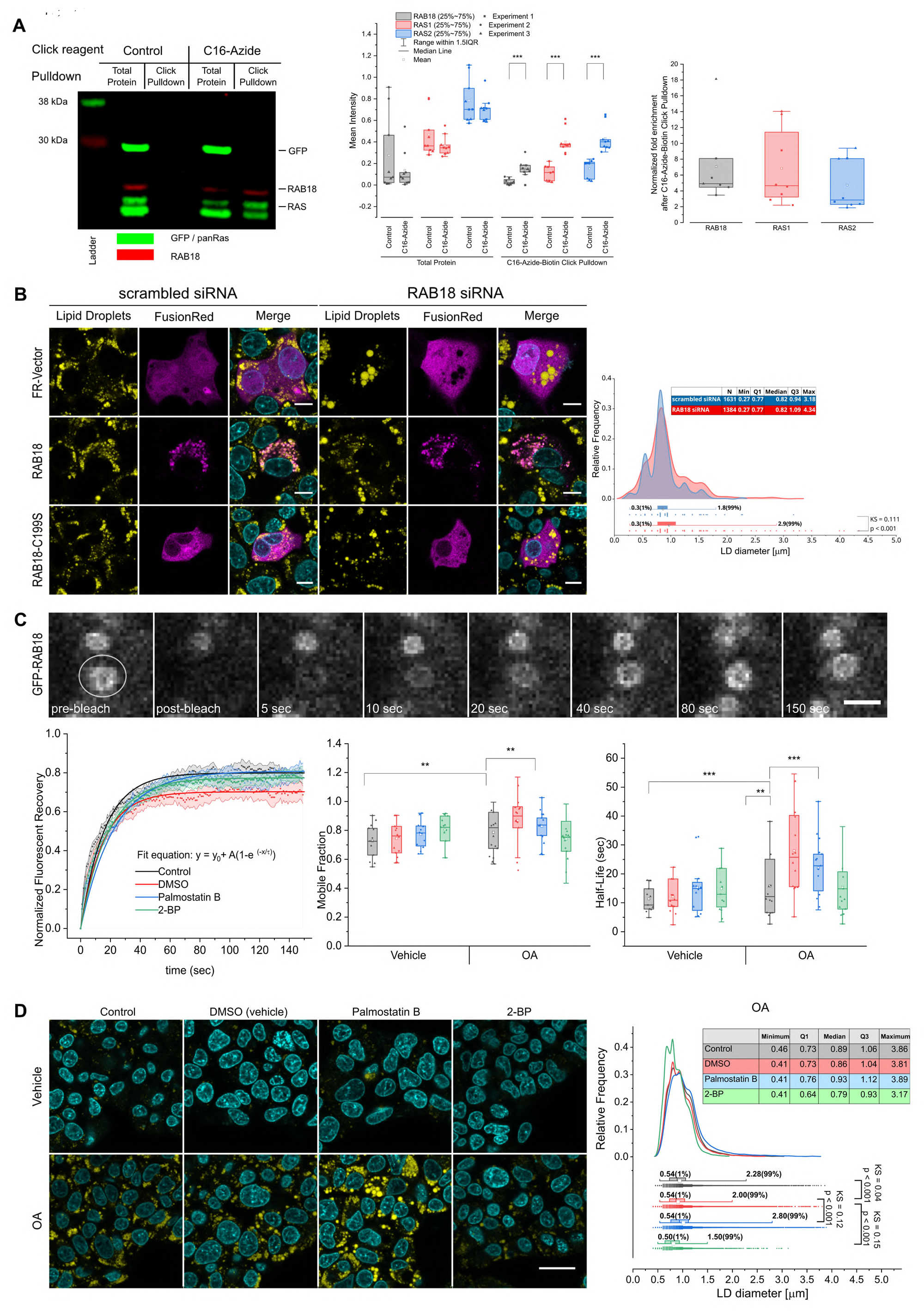
RAB18 localization is dependent on reversible C-terminal palmitoylation at Cys199. **A.** Representative fluorescent dual-color western blot of palmitoylated protein segregated from unpalmitoylated protein by Click-chemistry based biotin pulldown. RAS (green, ∼21 kDa) and RAB18 (red) were detected in the total protein as well as the palmitate-azide click reaction, whilst ectopically expressed GFP (green, ∼27 kDa) was not detected in the pulldown. Mean intensity of bands was quantified and normalized to total protein input and pulldown without palmitate-azide incubation. RAB18 and Ras were enriched in the pulldown by feeding palmitate-azide. number of experiments (N), Number of pulldowns (n): n, N_control,Click-Pulldown_= 8, 3; n, N_Control,Total-protein; C16-Azid, Click-Pulldown; C16-Azide, Click-Pulldown_=9, 3; statistical significance was tested with the Two-sample Kolmogorov-Smirnov test. **B.** Representative images of HepG2 cells expressing palmitoylation deficient RAB18-C199S. RAB18-C199S mis-localized and failed to rescue the LD size in cells with RAB18 depletion. Number of experiments (N) and Total number of LDs quantified (n): n, N_C199S-scrambled_=1631, 3; n, N_C199S-siRNA_=1384, 3; Scalebar: 10 µm. **C.** Representative timeseries and recovery graphs of the fluorescent recovery of GFP-RAB18. An increase of mobile fraction and localization half-life could be detected after LD formation. Mobile fraction and localization half-life were not affected by the inhibition of de-palmitoylation (Palmostatin B) but reduced by the inhibition of palmitoylation (2-Bromopalmitate) after LD formation. Number of FRAP curves (n) number of experiments (N): n, N_Vehicle,Control_=13, 3; n, N_Vehicle,DMSO_=18, 4; n, N_Vehicle, PalmostatinB_=12, 3; N_Vehicle, 2-Bromopalmitate_=19, 4; N_OA, Control_=18, 3; N_OA, DMSO_=24, 4; N_OA, PalmostatinB_=17, 3; N_OA, 2-Bromopalmitate_=25, 4. **: p<0.01; ***:p<0.001, two-sample t-test. Error-bars represent standard error Scalebar: 2µm **D.** Representative images of cells treated with palmitoylation-cycle inhibitors prior to 24h induction of LD formation with OA (left) as well as the LD size KDE after 24h of LD formation (right panel). Inhibition of depalmitoylation resulted in larger, inhibition of palmitoylation in smaller LDs. Number of experiments (N) and Total number of LDs quantified (n): n,N_Control_=31650, 4; n, N_DMSO_=61533,6; n, N_PalmostatinB_=33528, 3; n, N_2−Bromopalmitate_=32331, 3; Scalebar: 25µm

To zero-in on the exact site of palmitoylation on RAB18, we ectopically expressed GFP, GFP-RAB18 or the GFP-Rab19C199S mutant lacking the postulated palmitoylation site. Western blot analysis showed that GFP-RAB18 is detected in the palmitoylated protein fraction while GFP-RAB18C199S and GFP alone is not (**Fig. S2**). This confirms that the site of S-palmitoylation in RAB18 is Cys199 in its HVR.

To investigate if localization of RAB18 through S-palmitoylation is important for its function to constrain lipid droplet size, we performed a rescue experiment with ectopically expressed unpalmitoylatable FusionRed-RAB18C199S in cells with endogenous RAB18 depletion. Indeed, the unpalmitoylatable mutant neither localized to lipid droplets nor was it able to fully rescue the LD-enlargement caused by RAB18 depletion **(Fig 3B)**. These results further underscore the importance of palmitoylation in the localization and thereby the function of RAB18 lipid droplets.

Since S-palmitoylation is chemically reversible, and ubiquitously expressed thioesterases perform cell-wide depalmitoylation, wild type RAB18 should be continuously depalmitoylated on the lipid droplet membrane, forming the so-called ‘acylation cycle’. Fluorescence recovery after photobleaching (FRAP) of GFP-RAB18 individual entire lipid droplets showed that RAB18 is continuously turned-over on the LD membrane with a half-life of approximately ∼10s. Thus, RAB18 localization to the lipid droplet is dynamic. Perturbation of the acylation cycle by either thioesterase inhibition by Palmostatin B, or by inhibition of palmitoylation by 2-Bromopalmitate altered the half-life of RAB18 on the lipid droplet under exogenous lipid load (**Fig 3C**). These perturbations of the acylation cycle also led to changes in the function of RAB18 to constrain lipid droplet size. Specifically, inhibition of depalmitoylation by Palmostatin B led to an increase in droplet size, while inhibition of palmitoylation by 2-Bromopalmitate led to a decreased lipid droplet size under exogenous lipid load (**Fig 3D**).

However, inhibitors affecting the acylation cycle almost certainly affect the acylation cycles of numerous S-palmitoylated proteins, while 2-Bromopalmitate is also a known inhibitor of lipid synthesis. While this experiment is informative, pleiotropic effects of these inhibitors cannot be ruled out. We therefore investigated the cellular processes that RAB18 affects when it is localized to the lipid droplet membrane.

### RAB18-depletion induced LD-enlargement is lipophagy-dependent and lipolysis-independent

The size of LDs is maintained by a balance of anabolic and catabolic processes. LD formation and initial enlargement is known to occur on the ER and involve RAB18^7^. However, since the depletion of RAB18 on **LDs** (not the ER) leads to their enlargement, we hypothesized that RAB18 localized to the ER may be affecting a catabolic process instead. The two catabolic processes that affect LDs are lipolysis mediated by the triglyceride lipase ATGL, and ‘phagic’ degradation by autophagosomes^17^. In hepatocytes in particular, an unusual form of direct phagic degradation of LDs by fusion with lysosomes is reported^18^. We therefore investigated if either lipolysis or lipophagy plays a role in generating the enlargement of LDs seen under RAB18-depletion conditions.

HepG2 cells were treated with selective lipase inhibitors - ATGListatin, and inhibitor of lipolysis^19^ or Lalistat 2, an inhibitor of the lysosomal acid lipase responsible for lysosome mediated degradation^20^. As before, the depletion of endogenous RAB18 led to an increase in average lipid droplet size, but inhibition of lipolysis by ATGListatin neither enhanced nor retarded the enlargement of LDs **(Fig. 4A)**. In contrast, inhibition of lipophagic degradation by Lalistat 2 effectively reversed the RAB18-depletion induced LD enlargement providing a clue that RAB18 on the LD affects their lipophagy. Further, Lalistat 2 also substantially increased the *number of LDs per cell* indicating that LD-lipophagy was attenuated through inhibition of lysosomal activity.

**Figure 4.**
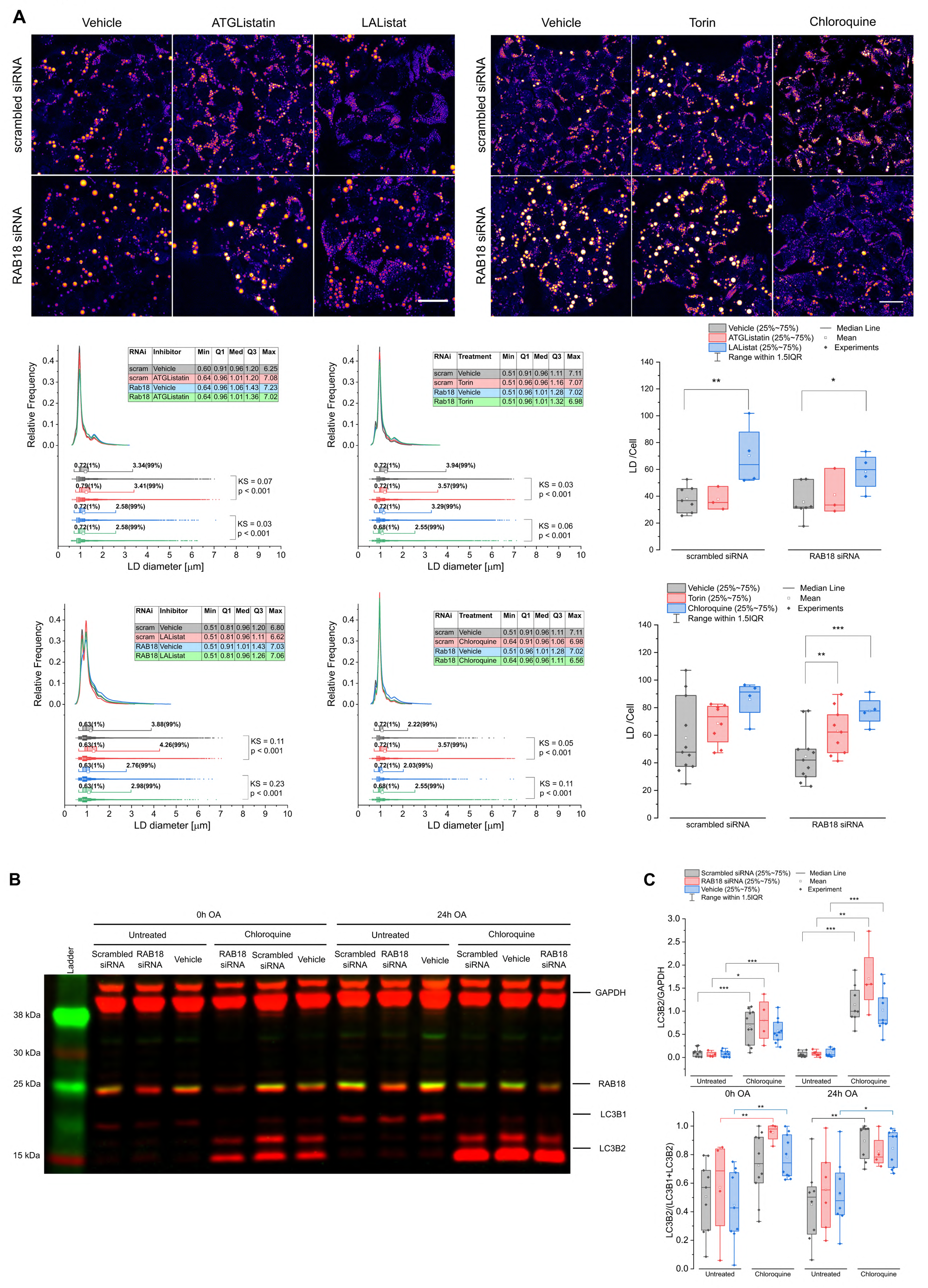
RAB18 depletion phenotype can be reverted by autophagy inhibition but does not affect LC3B levels. **A.** Representative images of HepG2 cells with RAB18 depletion after 24h of LD formations and inhibition of lipolysis (ATGListatin), lipophagy (Lalistat 22) as well as autophagy enhancement (torin2) and inhibition (chloroquine). LDs were detected with FCARS via C-H bond stretches at 2847 cm^-1^. LD size and number was not affected by lipolysis inhibition, but reduced by lipophagy inhibition. Enhancing autophagy did not affect the LD size distribution, but increased LD number in cells with RAB18 depletion. Inhibition of autophagy reverted the RAB18 depletion phenotype and doubled the LD number per cell. Number of experiments (N) and Total number of LDs quantified (n): n, N_scrambled siRNA, Vehicle ATGListatin/Lalistat 2_=291535, 7; N_scrambled siRNA, ATGListatin_= 246524, 3; n, N_scrambled siRNA, Lalistat 22_= 149859,4; n, N_siRNA RAB18, Vehicle, ATGListatin/Lalistat 2_ = 277484, 7; n, N_siRNA RAB18, ATGListatin_= 217353, 3; n, N _siRNA RAB18, Lalistat 22_ = 314481,4; n, N_scrambled siRNA, Vehicle, torin/Chloroqune_= 524388, 11; n,N_scrambled siRNA, torin_= 432252, 8; n, N_scrambled siRNA, chloroquine_= 169771,4; n, N_siRNA RAB18, Vehicle, torin/Chloroqune_ = 422650, 11; n, N_siRNA RAB18, torin_= 375945, 9; n N_siRNA RAB18, chloroquine_ = 182110,4. Two-sample T-test. **B.** Representative fluorescent dual-color western blot detecting LC3B, RAB18 and GAPDH in HepG2 with autophagy inhibition analogous to the experiment shown in A. Protein levels were analyzed prior and post induction of LD formation. **C.** Quantification of LC3B2 levels and ratio of LC3B2 to total LC3B. Inhibition of autophagy resulted in a significant increase of LC3B2 protein levels, but no alterations in LC3B2 level could be detected in cells with RAB18 depletion. N_scrambled siRNA, 0h, untreated_=9; N _RAB18 siRNA, 0h, untreated_=4; N _Vehicle, 0h, untreated_=9; N_scrambled siRNA, 0h, chloroquine_=10; N_RAB18 siRNA, 0h, chloroquine_=4; N_Vehicle, 0h, chloroquine_=10; N_scrambled siRNA, 24h, untreated_=8; N_RAB18 siRNA, 24h, untreated_=6; N_Vehicle, 24h, untreated_=8; N_scrambled siRNA, 24h, Chloroqune_=8; N_RAB18, siRNA, 24h, chloroquine_=4; N_Vehicle, 24h, chloroquine_=9; Two-sample Kolmogorov-Smirnov test. *: p<0.05; **: p<0.01; ***: p<0.001. Scalebar: 25 µm;

We therefore tested the effect of RAB18 depletion on lipophagy using further pharmacological perturbations through treatment with the autophagy enhancer torin as well as the global prevention of fusions of LDs with autophagosomes/lysosomes using chloroquine. In line with previous results, enhancement of autophagy with torin increased lipid droplet size in HepG2 cells, with the effect even more pronounced under RAB18 depletion. Conversely, treatment with chloroquine reduced the lipid droplet size to near control levels in HepG2 with endogenous RAB18 depletion and increased the number of LDs observed per cell. These data indicated that RAB18 depletion affected the autophagy/lysolipophagy of lipid droplets and resulted in an increase in LD size.

As the effects of RAB18 depletion on LD size and number could clearly be reversed by modulating autophagy/lipophagy, we investigated if RAB18 directly affects the autophagic flux in the cell. During autophagosome maturation, LC3B1 is converted to LC3B2, which is then degraded as autophagosomes fuse with lysosomes. RAB18 did not affect expression of LC3B2, LC3B1 nor the autophagic flux ratio. Similarly, the exogenous lipid load by OA did not show any influence on the autophagic flux. However, chloroquine caused a massive increase in LC3B2, which is consistent with previous reports that chloroquine blocks autophagic degradation (**Fig. 4B,C)**. In addition, chloroquine decreased LC3B1 and increased the ratio of LC3B2 to Total LC3B (LC3B2 + LC3B1), which has previously been used as an indicator of autophagic flux^21^. However, this interpretation should be treated with caution since the accumulation of LC3B2 may reduce LC3B1 expression as previously reported^22^. Nonetheless, together these data indicated that RAB18 does not affect the autophagic flux directly, while chloroquine did prevent phagic fusions globally as expected.

This leaves us with a conundrum – inhibition of a *degradative process* – lysolipophagy by chloroquine, leads to a *decrease* of lipid droplet size, along with a concomitant *increase in the number of droplets* (**Fig. 4A**), completely reversing the LD-enlargement caused by RAB18-depletion. Having eliminated lipolysis as well as any direct effects on the autophagic machinery, we then postulated that RAB18 *on the lipid droplet* must be instead modulating interactions of lipid droplets with phagic organelles. Such a modulation has been previously suggested^17^, but its effects on relevant cellular processes LD-dynamics remain unknown.

### RAB18 reduces LD-lysosome interactions to prevent LD enlargement and maintain LD numerosity

The ‘steady-state’ distribution of lipid droplet size as shown so far does not allow investigation of the effect of RAB18 on dynamic lipid droplet interactions with autophagic organelles. Instead, we used time lapse monitoring of lipid droplet interactions with autophagosomes and lysosomes under RAB18-depletion conditions and exogenous lipid load. Lipid droplets were visualized by BODIPY, autophagosomes by ectopic expression of CFP-LC3B and lysosomes by Lysotracker Deep Red. Interactions of lipid droplets with lysosomes (**Fig. 5A, Supplementary movie M3**). were 2-fold more frequent than with autophagosomes (**Fig. S5, Supplementary movie M4**). Importantly, RAB18 depletion increased interactions of LDs with lysosomes, with exogenous lipid load. As before, lipid droplet size was enlarged under RAB18 depletion conditions. Interestingly, the number of lipid droplets increases upon exposure to exogenous lipid load, but under RAB18-depletion conditions, this increase in lipid droplet number is dampened. In contrast, RAB18 depletion did not influence interactions with autophagosomes. This is in line with the minimal effect of autophagy modulation and rather underscores the importance of direct lysolipohagy of LDs^18^ in hepatocytes.

**Figure 5.**
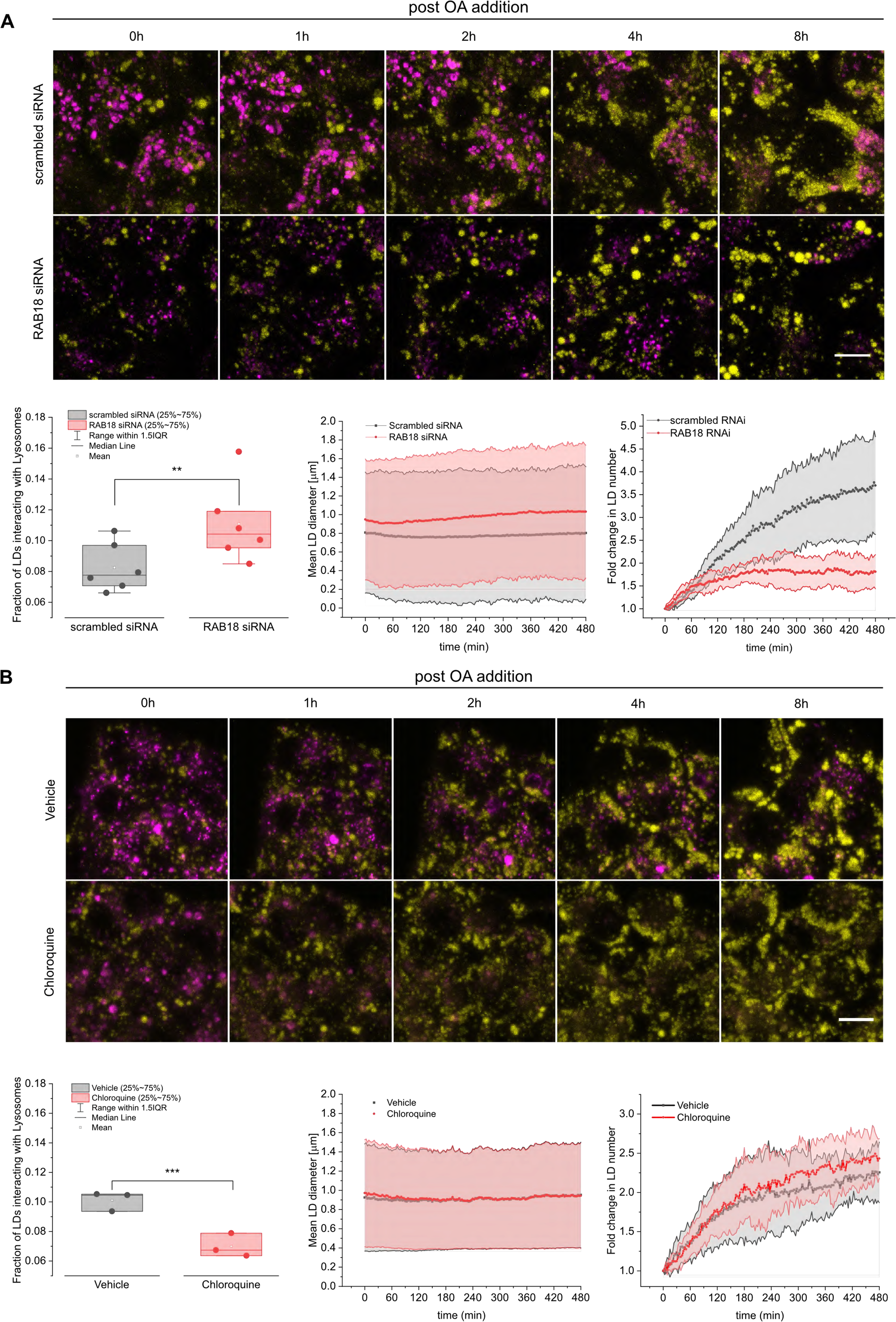
RAB18 depletion leads to a higher fraction of LDs interacting with lysosomes. **A.** Representative timeseries of the first 8h of LD formation after OA addition in RAB18-depleted cells. LDs were detected with BODIPY (yellow), lysosomes with lysotracker deep red (magenta). An increased fraction of LD was detected to interact with lysosomes in cells with RAB18 depletion. LD mean area was increased in cells with RAB18 depletion, and the number of newly formed LDs was decreased. Number of experiments (N), number of timeseries analyzed (n): n, N = 6, 3. **B**. Representative images of a timeseries of the first 8h of LD formation in cells pretreated with chloroquine. Pretreatment with chloroquine led to a reduction in the fraction of LDs interacting with lysosomes. There was no change in LD mean area whilst the number of newly formed LDs was slightly increased after 8h. Number of experiments (N), number of timeseries analyzed (n): n, N= 3, 3; Two-sample T-test. *: p<0.05 **: p<0.01. Scalebar: 10 µm

We have previously shown that chloroquine is able to reverse the LD-enlargement caused by RAB18 depletion. We therefore asked if this is due to a corresponding but ***reciprocal*** effect on interactions of LDs with lysosomes. As expected, chloroquine reduced the number of interactions of lipid droplets with lysosomes, which was accompanied by an increase in lipid droplet numbers (**Fig. 5B, Supplementary movie M5**).

These data reveal a general pattern wherein lipid droplet size and number are inversely related. RAB18 and chloroquine both prevent interactions of LDs with lysosomes, leading to more numerous but smaller lipid droplets.

### Pharmacological modulation of lipophagy prevents liver steatosis in primary hepatocytes and mice

It is apparent that lysolipophagy plays a prominent role in the determination of lipid droplet size and numbers. However, these conclusions were obtained from data in HepG2 cells. It is unknown if lysolipophagy is similarly important in lipid droplet dynamics in more native systems. If these processes are indeed similar in primary liver cells, the use of lysolipophagy inhibitors to prevent LD enlargement in steatotic liver disease is attractive for its therapeutic potential. We therefore investigated the effect of chloroquine to prevent LD-enlargement in primary human hepatocytes under exogenous lipid load. Treatment of primary human hepatocytes with chloroquine led to a concentration-dependent reduction in LD enlargement and corresponding increase in LD number **(Fig. 6)**. It is interesting to note that chloroquine nearly completely abolished LD-enlargement at 100 µM in PHHs.

**Figure 6.**
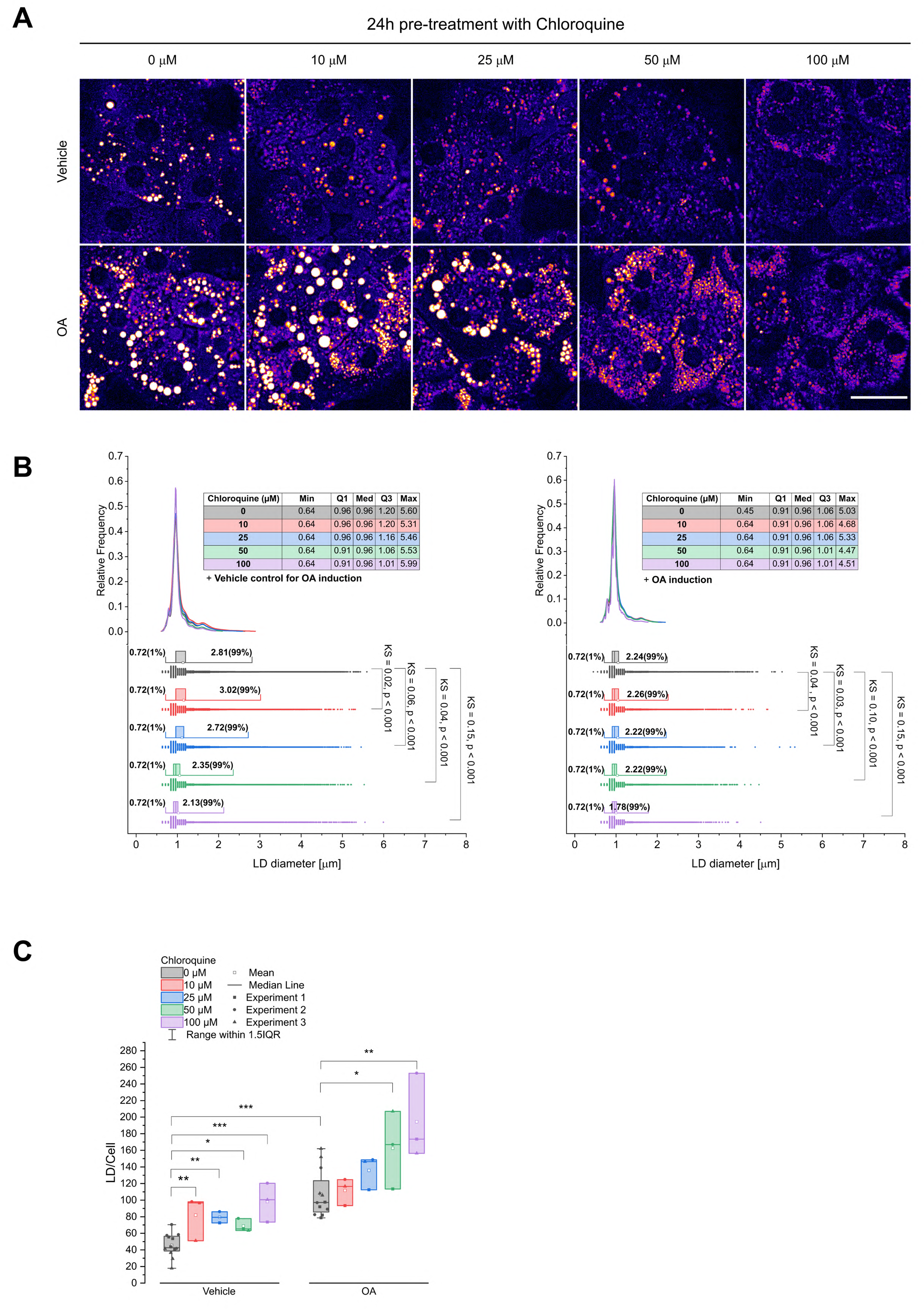
chloroquine treatment reduces LD growth in human hepatocytes. Representative images of primary human hepatocytes treated with increasing concentrations of chloroquine after 24h induction of LD formation with OA. LDs were detected with FCARS via C-H bond stretches at 2847cm^-1^. chloroquine treatment resulted in concentration dependent decrease n in LD size and increase in LD number. Number of experiments (N) and Total number of LDs quantified (n): n, N_Vehicle,0µM_ =152080, 3; n, N_Vehicle,10µM_ = 55342, 3; n, N_Vehicle,25µM_ = 39070, 2; n, N_Vehicle,50µM_ = 50519, 3; n, N_Vehicle,100µM_ = 103202, 3; n, N_OA,0µM_ = 398336, 3; n, N_OA,10µM_ = 87185, 3; n, N_OA,25µM_ = 105630,3; n, N_OA,50µM_ = 122286, 3; n, N_OA,100µM_ = 189104, 3;Two-sample T-test *: p<0.05; **: p<0.01; ***: p<0.001. Scalebar:25 µm.

Encouraged by these results, and leveraging the fact that chloroquine is a clinically approved drug, we performed a small-scale mouse study. Mice were put on a chloroquine regimen (60 mg/kg) and subsequently on a steatogenic diet. Intravital CARS imaging was performed every week on the livers of anaesthetized mice over 4 weeks to track the development of LDs (**Fig. 7A**). CARS leverages excitation of the C-H bond vibrations in lipids and detects them with much higher sensitivity than conventional histology on tissue sections. Quantification of LD size showed that mice on control diet had the lowest number and mean LD size (4100/mm^2^, 2.1 µm) for large lipid droplets. Mice on steatogenic diet showed more numerous and enlarged large LDs. (21125/mm^2^, 2.8 µm), while mice on steatogenic diet treated with chloroquine had significantly smaller lipid droplets (14317/mm^2^-,2.1 µm). Thus, inhibition of lysolipophagy prevented the formation of large LDs in vivo, effectively attenuating the development of steatosis (**Fig. 7B**). Due to mouse positioning on the imaging stage, it is only possible to obtain epi-CARS (E-CARS) signal in live mice. The E-CARS signal is produced by backscattering of the forward CARS (F-CARS) signal by large scatterers^23^, thereby biasing the E-CARS detection towards lipid droplets that are larger than or comparable to the wavelength of excitation light. We therefore also imaged frozen tissue sections from these mice, where the unbiased F-CARS signal could also be measured (**Fig. S7**). As expected, the F-CARS image shows the presence of much smaller lipid droplets that are detected in the E-CARS image and confirms that chloroquine treatment leads to a reduction of lipid droplet size in both control and steatogenic diet groups.

**Figure 7.**
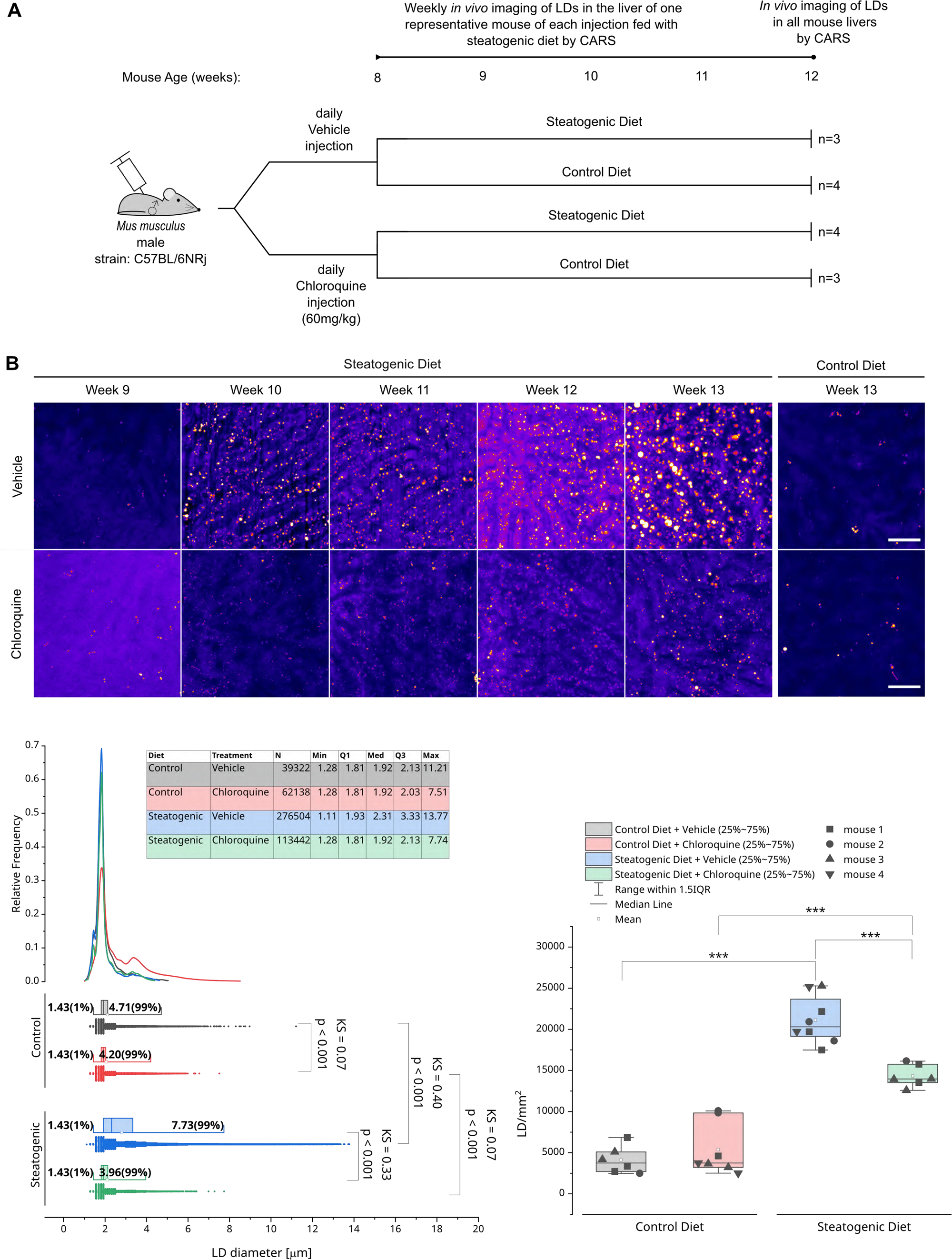
chloroquine treatment reduces LD growth in mice. **A.** Schematic overview of the conducted mouse experiment. At the age of 8 weeks, two populations of mice were injected daily with either 60mg/kg bodyweight chloroquine. At week 9 half of both populations were additionally put on a steatogenic diet. The LD growth in the steatogenic diet was monitored by in vivo imaging the liver LDs of one exemplary mouse of each condition. At week 13 all mice livers were imaged. **B.** Representative intravital ECARS images of murine livers, showing the onset of steatosis. LDs were detected via C-H bond stretches at 2847cm^•-1^. chloroquine treatment prevented the increase of LD size and reduced the number of LDs in mice on the steatogenic diet. Number of experiments (N) and Total number of LDs quantified (n): n, N_ControlDiet,Vehicle_ =39322,3; n,N_ControlDiet, chloroquine_= 62138,4; n, N _SteatogenicDiet, Vehicle_ = 276504, 4, n, N_Steatogenic Diet chloroquine_=136511, 3; Two-sample Kolmogorov Smirnov test. *: p<0.05; **: p<0.01; ***: p<0.001. Scalebar:50 µm

## Conclusion and Discussion

In this work, we establish that RAB18 modulates LD size and number through regulating their interactions with lysosomes. When LD-localized RAB18 is reduced, interactions between lipid droplets and lysosomes increase, resulting in a phenotype with lower numbers and larger size of LDs (Figure 6A,B). We show that RAB18 localizes to the LD membrane via post-translational S-palmitoylation and is dynamically turned over as part of its acylation cycle. Localization to LDs via the acylation is dependent on the GTP-loading state of RAB18, as is typically seen for Rab-family members^12,24^. Perturbation of the acylation cycle by thioesterase can replicate the phenotype of RAB18 depletion on the LD.

The inverse relationship between LD size and number is consistently observed during several types of interventions – RNAi for RAB18, pharmacological inhibition of LD-lysosome interaction, and inhibition of lysosomal lipophagy. Still, it appears paradoxical that inhibition of LD-degradation (a catabolic process) through lysolipophagy leads to smaller sizes of lipid droplets. This paradox can be resolved through a conceptual model which we term the ‘container-shortage’ concept. In this concept, it is assumed that the lipid load incumbent upon the cell must be stored in lipid droplets. RAB18 protects lipid droplets from lysosomal degradation by reducing their interactions with lysosomes. Upon RAB18 depletion, lipid droplets are degraded due to increased interaction with lysosomes, leaving fewer lipid droplets available to absorb the incumbent lipid load. Since fewer droplets must take up the same incumbent lipid load, the size of each individual droplet is inevitably larger. The container-shortage concept predicts that LD enlargement can be prevented simply by ensuring that a sufficient number of lipid droplets are present in the cell to absorb the incumbent lipid load. Indeed, pharmacological prevention of the degradation of lipid droplets by preventing their interaction with lysosomes (chloroquine) or by inhibiting of lysosomal activity (LaliStat2) also prevents LD enlargement.

RAB18 on the ER has been previously reported to be involved in the biogenesis of lipid droplets^5^. Therefore, it is possible that lower numbers of lipid droplets observed under RAB18-depletion reflect reduced biogenesis. However, prevention of LD degradation by chloroquine completely reversed the LD-enlargement seen under RAB18 depletion – indicating that the degradation of lipid droplets is the more potent factor in determining LD numbers and thus LD size.

Lipid droplets begin life with the formation of a TAG enriched ‘lens’ on the ER membrane. This lens detaches to form a vesicle with diameter between 80-200 nm, which is called a nascent lipid droplet^7^. The growth of nascent lipid droplet to super-droplets seen in disease conditions seems to occur in at least 2 phases. In Phase 1, nascent lipid droplets grow to approximately 5 µm in diameter, possess RAB18 on the membrane and are subject to the control processes described above. In Phase 2 these lipid droplets further fuse with each other via the action of Perilipin 1/FSP27 to form a single unilocular super-droplets. It should be considered that the present work focuses on regulation of LD size in Phase 1, which is clinically described as ‘microsteatosis’ and precedes and enables progression to macrosteatosis (Phase 2)^25^.

The relevance of the above findings to clinical observations in MAFLD are underscored by the efficacy of chloroquine to prevent the development of microsteatosis in a mouse model of steatogenic diet. chloroquine, which prevents LD-lysosome fusions also prevented enlargement of lipid droplets in mouse livers upto 4 weeks. As chloroquine is a clinically approved drug, the potential for therapeutic application is attractive, with the caveat that as MALFD progresses further complexity may be introduced. For example, while cell-based experiments showed that the lipid droplet size and number were generally anti-correlated irrespective of the cause, this did not seem to be the case in the mice. On the contrary, in mice, inhibition of autophagy caused a reduction of lipid droplet size *and* number. While serendipitous in terms of preventing the development of steatosis, this observation is not predicted by the container shortage model. Whether this can be attributed to additional complex processes in living mice compared to cultured cells, or the different time periods (4 weeks vs. 24 hours) or different lipid delivery (complex diet vs. oleic acid suspension) remains to be elucidated.

In conclusion, we characterize the localization mechanism of RAB18 to the lipid droplet membrane, demonstrate its function in preventing lipid droplet degradation and develop the container-shortage model, providing a rational framework linking cellular processes underlying lipid droplet dynamics to the pathogenesis of steatotic disease.

## Supplementary Information

### Methods

#### Materials

Material as well as media used in this publication are listed in Table S3 and Table S4 respectively. Primer sequences are detailed in table S1.

#### Cell line

Human [*Homo* sapiens] liver carcinoma cells (HepG2, ATCC) were cultured in HepG2 cell medium (See S4) and kept at 37°C and 4% CO_2._ Cells werepassaged twice a week until passage 27.

#### Generation of fluorescent tagged RAB18 variants

RAB18-ORF was amplified from a Turbo-GFP-ORF vector (Origene) and inserted into FusionRed-and GFP2-C-tag expression vectors (Evrogene). The fluorescently tagged *rab18* was mutated (for exact sequence changes table S2) using a site directed mutagenesis kit (Agilent). Sec61b-ORF was obtained from the BFP-SEC61 beta plasmid gifted by Gia Voeltz to be inserted downstream of the RAB18-ORF.

#### HepG2 Plasmid transfection

Overexpression plasmids were transfected using the Effectene kit (Qiagen). For each well of an 8-Well chambered slide (Ibidi), 1µg plasmid was added to 100µl EC Buffer. 8µl Enhancer reagent per µg plasmid was added and the solution quickly vortexed (2-5 seconds). Per well 1.25µl of Effectene reagent was added, before vortexing the solution again. The transfection mix was incubated for 5-10 minutes at room temperature and added to the cells. 6-well plates were treated analogous using 2µg of plasmid and 5µl Effectene in 200µl EC per well.

#### HepG2 inverse RNA transfection

For each well of an 8-Well chambered slide 4.8-9.6 pmol scrambled or RAB18 targeting siRNA (Origene) were added to 40µl of DMEM medium without supplements (PAN-biolabs). After addition of 0.4µl RNAimax (Sigma-Aldrich) the mix was added to the wells and incubated for 25-30 minutes at room temperature. 260µl of cell suspension was added on the siRNA transfection mix. 6-well plates were treated analogous using 500µl of DMEM, 30 pmol of siRNA, 5µl RNAimax and 2ml of cell suspension per well.

#### Characterization of RAB18 depletion phenotype

HepG2 cells were seeded in an 8-well chambered slide and transfected with siRNA. 72h post siRNA transfection LD were stained with 1µM BODIPY (Invitrogen) for 1h. The slide was equilibrated on a confocal microscope for 30 minutes at 37°C and 5% CO_2_, before 200µM BSA complexed oleic Acid (Sigma Aldrich; abbr. OA) with 1µM BODIPY (Invitrogen) were added to observe LD formation. The LD formation was imaged at 3 positions in each well in 15 minutes steps over 20h using a 40х objective (Zeiss, LSM880).

#### Rescue of RAB18 depletion phenotype

HepG2 cells were seeded in an 8-well chambered slide and transfected with siRNA. After 8h-o.n. incubation cells were transfected with FusionRed tagged RAB18 variants. 72h post siRNA transfection LD growth was induced by addition of 400µM OA to the cells for 20h. The next day, cells were stained with 0.1µM BODIPY for 45 minutes followed by 15 minutes of nuclei staining with 1µg/µl Hoechst 33342 (Invitrogen). HepG2 cells were washed thrice with prewarmed medium, before taking 5х5 tile-scans of each well using a 63х objective (Leica, SP8). Single transfected cells of each tile-scan were picked and LD size manually quantified.

#### FRAP analysis of RAB18 localization

HepG2 were seeded in an 8-well chambered slides and transfected with GFP-RAB18. Transfected cells were treated with 400µM OA, BSA as vehicle control or left untreated for 20h. Palmitoylation was inhibited by incubating the cells 50µM 2-Bromo-palmitate (Sigma-Aldrich) for 4 hours. De-palmitoylation was inhibited by incubation with 10µM Palmostatin B (Sigma-Aldrich) for 10 minutes. DMSO (Sigma-Aldrich) was used as a solvent control. LD localized GFP-RAB18 was imaged for 5 seconds before bleaching it for 5 seconds with an Argon laser. Post-bleach imaging was conducted each second for 150 seconds using a 63х objective (Leica SP8). Quantification was done via a custom-made FIJI macro. The mean intensity of the bleached area was corrected for imaging dependent photobleaching and double normalized to an unbleached reference^1^. The bleach corrected data was subtracted by the minimum value and normalized to the maximum. Each curve was manually checked for artifacts. A global mono exponential fit was performed on the normalized FRAP curves to calculate the mobile fraction A and the time constant τ using OriginPro.

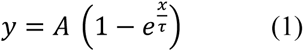

Localization half-life was calculated based on the obtained τ values (2).

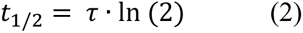

#### LD size modulation by palmitoylation manipulation

HepG2 cells were seeded in 8-well chambered slide. 20h post seeding the cells were treated with 50µM 2-Bromopalmitate and 100µM PalmostatinB for 4h. After the incubation, cells were treated with 400µM OA, BSA as vehicle control and DMEM medium for 24h. LD and nuclei were stained using BODIPY and Hoechst respectively. Stained cells were imaged in 5х5 tile scans using a 63х objective (Leica, SP8). The tile-scans were stitched and LD area was quantified using StarDist algorithm.

#### Test of lipolysis and autophagy regulating inhibitors

HepG2 cells were seeded in 8-well chambered slide and transfected with siRNA. 48h post siRNA transfection lipolysis was inhibited with 50µM ATGListatin(Sigma-Aldrich), lipo-or autophagy were inhibited with either 50µM LAListat2 (Sigma-Aldrich) or 30µM Chloroquine (Sigma-Aldrich) respectively. Autophagy was activated with 1µM Torin2 (Sigma-Aldrich). DMSO was used as vehicle control. After 24h incubation with the compounds, cells were incubated with 400µM OA for 24h. HepG2 cells were subsequently imaged with CARS at a wavenumber of 2847cm^-1^ using an IR-optimized 40х objective (Leica, SP8). 5х5 tilescans were taken, stitched and background was removed using the rolling ball background subtraction FIJI plugin with a width of 10 pixel. LDs were segmented using the StarDist algorithm and a custom trained model. Cell number was quantified manually.

#### Palmitoylated protein isolation by Azide-palmitate pulldown

HepG2 cells were seeded in a 6-well plate (Sarstedt). Two wells were each transfected with GFP, GFP-RAB18 or GFP-RAB18C199S 8h post seeding. Once confluent the medium was exchange for 1ml medium containing 500µM palmitic acid (Sigma-Aldrich) or palmitic acid azide (Invitrogen) and incubated for 5h. The cells were lysed in ice-cold RIPA buffer (Thermo Fisher Scientific) containing protease inhibitors (Roche) and 100µM PalmostatinB to prevent de-palmitoylation. Cell debris was removed by centrifuging the lysate at 10,000хg for 5 min at 4°C. 20µl were set aside as input control. To initiate the copper free click-reaction 52.5 nmol DBCO-biotin (Jena Bioscience) was added to the remaining protein solution and incubated at room temperature whilst shaking. The protein concentration in the input lysate was determined via BCA Assay (Thermo Fisher Scientific). 10 µg of the ‘input control’ protein were denatured by boiling at 95°C for 5 minutes in 20µl sample reduction buffer (4х Sample buffer, Licor; 10х Sample Reducing agent, NuPage) 100µl of streptavidin covered magnetic beads (Thermo Fisher Scientific) were washed three times in TBST. For storage the beads were finally resuspended in 100µl PBS. After 1.5 h the DBCO incubation beads were recaptured and PBS removed. From the DBCO-biotin reaction 300µg were added to the beads and the volume was adjusted to 200µl with RIPA buffer containing Palmostatin B. The beads were incubated for 30 minutes whilst shaking, before they were separated from the supernatant with a magnetic rack. The supernatant was removed and the beads were washed three times with PBS. Bound protein was eluted from the beads by boiling the beats in 20 µl sampled reduction buffer at 95°C for 5 minutes. The beads were separated from the denatured protein using the magnetic rack and the eluate analyzed via western blotting.

#### Macro-autophagy in cells with RAB18 depletion

HepG2 cells were seeded and transfected with siRNA in two 6-well plates. Cells were mock treated with the transfection reagent. 48h post transfection cells were treated with PBS or 20µM Chloroquine an incubated for 24h hours. The cells in one plate were lysed in ice cold RIPA buffer supplemented with protease inhibitors. 400µM OA was added to the remaining plate and the cells were incubated for additional 24h, before being lysed. Cell debris was removed by centrifuging the lysate at 10,000хg for 10 min at 4°C. 20µg of total protein were denatured by boiling at 95°C for 5 minutes in sample reduction buffer.

#### Western blotting

For analysis of LC3B and RAB18 by Western blotting, denatured protein was separated by electrophoresis on a pre-cast 4-12% Bis-Tris gradient gel (Invitrogen) at 200V in a mini-gel tank (Invitrogen) for 30 minutes. The protein was transferred onto a PVDF membrane for 35 minutes at 20V. Blots of palmitoylation pulldown were dried overnight at 4°C and rehydrated the next day. The membrane was blocked 1h at RT in TBS blocking buffer. Primary antibodies were diluted in 10 ml TBS blocking buffer as given in table S5. The blocking solution was exchanged for the primary antibody solution and incubated over night at 4°C. The fluorescent labeled secondary antibodies (Goat anti rabbit 680nm, Donkey anti mouse 800nm) were diluted 1:20000 in TBS blocking buffer. After overnight incubation the blot was washed in TBST and the secondary antibody solution was added for 2 hours at room temperature. The blot was imaged with a Licor Odyssey scanner with automatic illumination settings. The resolution was set at 169µm and highest quality.

##### Western blot quantification

Westernblots were quantified using FIJI. For palmitate pulldown assays ROIs were chosen for the bands of RAS1 and RAS2 as well as RAB18. The product of the protein bands’ mean intensity and area (integrated density) was measured and the background subtracted. The fold of enrichment through azide incubation was calculated for cells treated with and without palmitate-azide. To account for input variation the enrichment was normalized to the ratio of input protein with and without azide.

For LC3B level evaluation LC3B and RAB18 bands’ integrated densities were measured and normalized to the values of GAPDH. Cells transfected with siRNA but showing more than 10% of mock transfected cells’ RAB18 protein level were disregarded in the LC3B evaluation.

#### LD-Autophagosome interaction imaging

HepG2 cells were seeded in 8-well chambered slides and transfected with siRNA. 48h post siRNA transfection cells were transfected with a plasmid overexpressing fluorescent labelled LC3B-CFP encoded on pEX-CFP-hLC3WT which was a gift from Isei Tanida (Addgene plasmid # 24985; http://n2t.net/addgene:24985; RRID:Addgene_24985). 72h post siRNA transfection LDs were stained with 1µM BODIPY. LD formation was induced by addition of 400µM OA with 1µM BODIPY. Cells were imaged every 5 minutes for 4h on a Leica SP8 using a 63х objective. Focus was readjusted to counteract focus drift. Timeseries were concatenated and contrast maximized.

#### Lysolipophagy imaging

LD-lysosome interactions were imaged in HepG2 cells with and without RAB18 knockdown as well as HepG2 treated with 30µM chloroquine. LD were stained with 1µM BODIPY and lysosomes 25nM Lysotracker deep red for 20-30 minutes. LD formation was induced by adding 400µM OA with 1µM BODIPY. Z-stacks with 1µm stepsize were taken every 3 minutes o.n, using a 63х objective (Leica, SP8). For analysis stacks were maximum projected, bleach corrected and contrast maximized over the duration of the time series.

#### Autophagy and Lysolipophagy image analysis

Images of cells transfected with CFP-LC3B were first blurred and then thresholded to create a mask of transfected cells. This mask was applied to segment LDs in transfected cells. LD number and size were quantified for each experiment using the StarDist algorithm with a custom trained BODIPY model. Autophagosomes were segmented using the same model and labelled. LDs positive for at least one autophagosome label were counted as interacting LDs. Interacting LDs were normalized to total LD number to calculate the fraction of interacting LDs.

Image analysis of lysolipophagy was analyzed analogous without the initial segmentation of transfected cells.

#### Primary human hepatocyte (PHH) treatment with Chloroquine

Primary human hepatocytes (PHHs) taken from a female human donor with a BMI of 28.2 were seeded in 4-Well chambered slides coated with Rat Collagen (Roche). Upon thawing, the cells were suspended in 5 ml prewarmed PHH Medium (see S4). 200,000 cells were added to each well of the coated chamber and incubated for 3 hours at 37°C to attach. After attachment, the medium was aspirated thoroughly, and a second layer of collagen was applied. Fresh medium was added to the cells in sandwich culture. Primary human hepatocytes were kept at 37°C and 5% CO_2_ o.n.. The PHHs medium was exchanged with fresh WME supplemented 10µM, 25µM, 50µM or 100µM Chloroquine. PBS was used as vehicle control. The cells were incubated for 24h, before 400µM BSA complexed OA was added for 24h. LDs were detected with CARS imaging in 5х5 tilescans at 2857cm^-1^.

#### Treatment of mice on steatogenic diet

All experiments were approved by the official State animal care and use committee (LANUV, Recklinghausen, Germany AZ 84_02.04.2016.A473). From eight weeks-old male C57BL/6NRj mice with a body weight ranging between 20 to 25 grams. 4 groups of 7 mice each were formed. The mice were housed under 12 h light/dark cycles at controlled ambient temperature of 25°C with free access to water and to Ssniff R/M-H, 10 mm standard diet (Ssniff, Soest, Germany) respectively D16022301 Research diet (BROGAARDEN Smedevangen 5 350 Lynge, Denmark) Twenty-eight mice were divided in two groups. Group one was subjected to daily intra peritoneal injection with 60mg/kg body weight chloroquine phosphate solved in PBS. The control group two was injected with corresponding volume of PBS. The mice were treated this way for one week before the liver of one mouse of each group was imaged using intravital CARS imaging described below. After one week of chloroquine treatment seven mice of each group were set either on a steatogenic or a control diet. Mice were fed for four weeks with one representative mouse of chloroquine and untreated mice fed with research diet being imaged each week. After 4 weeks of diet all mice livers were imaged and subsequently euthanized. The livers were collected and fixed for 2 days in 4% PFA (Roth) at 4°C. At day 3 the solution was changed to 30% Sucrose (Sigma-Aldrich) and incubated for 3 days at 4°C. At day 5, the tissue was embedded in OCT (Fischer scientific) and stored at-80°C.

#### Intravital CARS imaging

Mice were anaesthetized with a combination of ketamine (64mg/kg) (CD-Pharma), xylazine (7,2 mg/kg) and acepromazine (1.7mg/kg) given intraperitoneally. The abdomen of the animal was shaved, and a ∼1.5 cm midline incision made to expose the xiphoid process which was retracted to allow dissection of the falciform ligament. The left lobe of the liver was gently exteriorized and the animal inverted onto a glass coverslip mounted within a custom-made imaging platform. The liver was covered with sterile PBS-soaked gauze to prevent dehydration. Images were acquired on a Leica SP8 in a light proof environmental chamber, which was maintained at 37°C and constant humidity. Both ECARS and FCARS as well as second harmonics imaging was conducted in 5х5 tile-scans taken at 2847cm^-1^. Tilescans were stitched, cropped to a rectangular area and a rolling ball background subtraction with a size of 40 pixels was performed using the FIJI plugin. LD were automatically segmented using the StarDist algorithm.

#### CARS imaging of cryoslides

Frozen tissue was cut with a cryotome in 5µm slides and transferred to a microscopy slide. Slides were kept at low temperatures for imaging. The frozen cuts were covered with a plasticine coverslip and imaged using CARS at 2800 cm^-1^. Three 5х5 tilescans were obtained per slide, stitched and analyzed using a custom StarDist model.

#### StarDist application for LD segmentation

The Jupyter notebooks supplied in the tutorials by the developer^2^ were followed to train a model for BODIPY stained and CARS detected LDs using hand annotated images as ground truth. The model was compared to test sets not used for training to verify correct LD number and area detection. The code in the notebooks was modified to allow batch processing of images saving resulting output as a ROIset. A custom-made FIJI macro was used to read out Area and number of LDs.

#### Statistical Analysis

For statistical analysis the OriginPro software was used. Datapoints outside the range of twice the standard deviation were labeled as outliers and excluded from the calculations. When normal distribution was not rejected by the Shapiro-Wilk test, the p-values were calculated using a two-tailed unpaired two-sample t-test and equivalent variance was tested for using the two-sample test for variance. When normality was rejected a two-tailed, two-sample Kolmogorov Smirnov test was used instead. Due to the large number of samples, ranging from n = 3000-250,000 lipid droplets, the conventional significance threshold of 0.05 is not useful. Instead, the Kolomogorov-Smirnov (KS) test statistic is reported as an indicator of effect size. The KS-statistic is 0 for two identical distributions and increases as the compared distributions become more disparate. The test used is given in the figure legend as well as the number of experimental replicates.

#### Data and Code availability

All data generated and analyzed are included in the manuscript and supporting files. Source data are provided with this paper. Models, macros and Jupyter notebooks used for the generation of the presented data are available under the following repository: https://cumulus.ifado.de/d/b588f4051a984fd38a19.

## Supplementary Tables

**Table S1:**
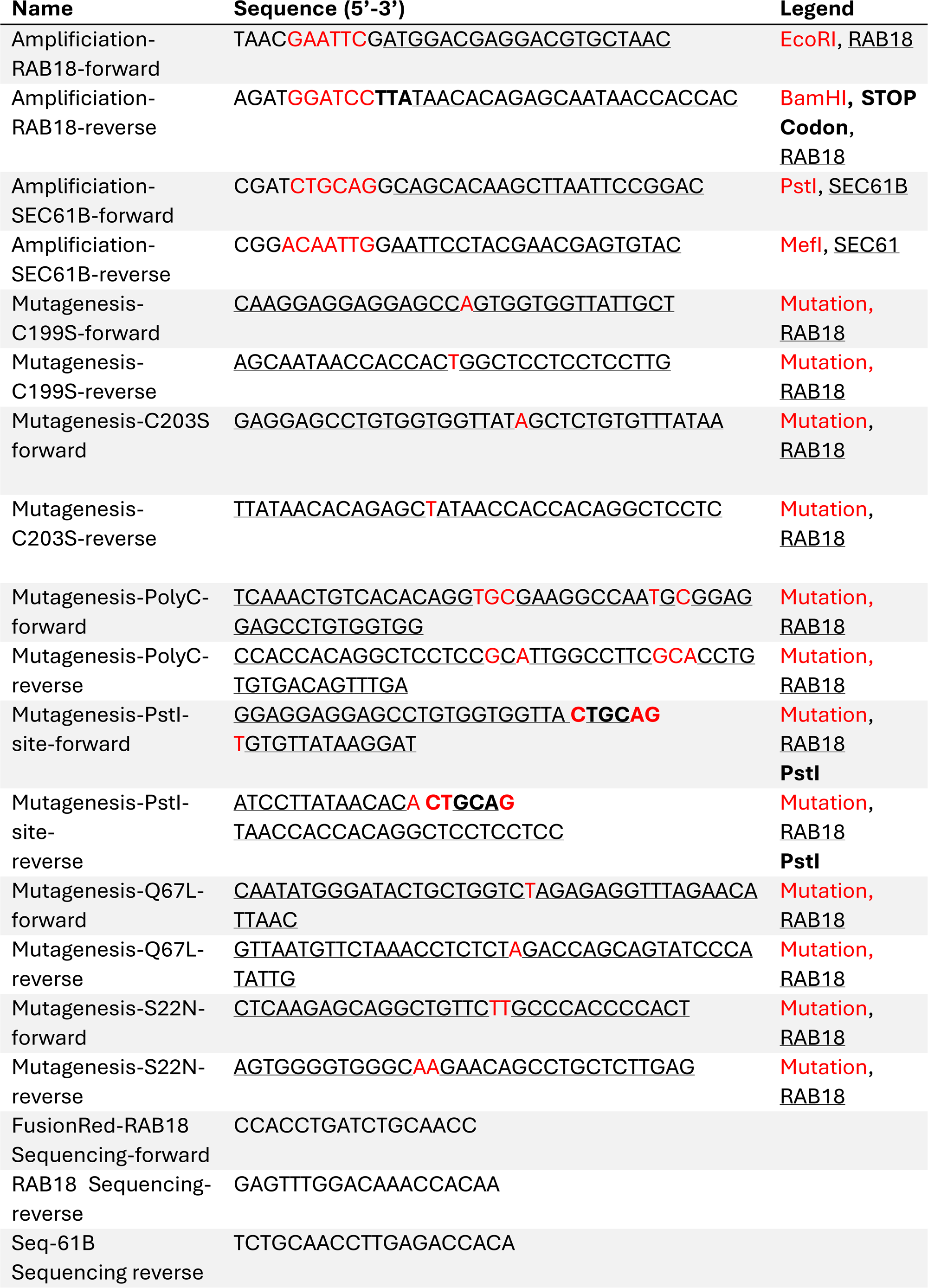
Primer

**Table S2:**
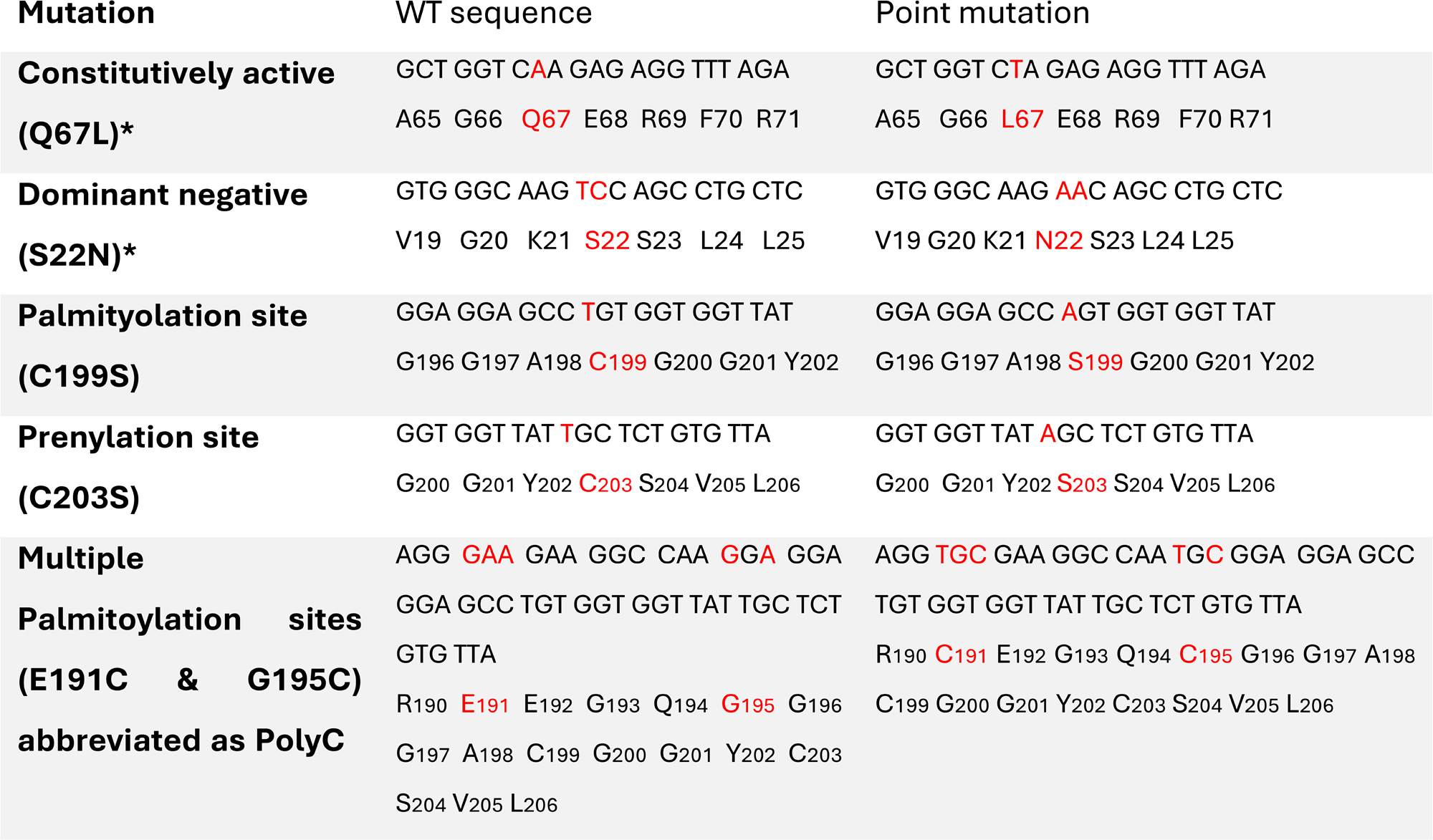
Detailed Mutation overview

**Table S4:**
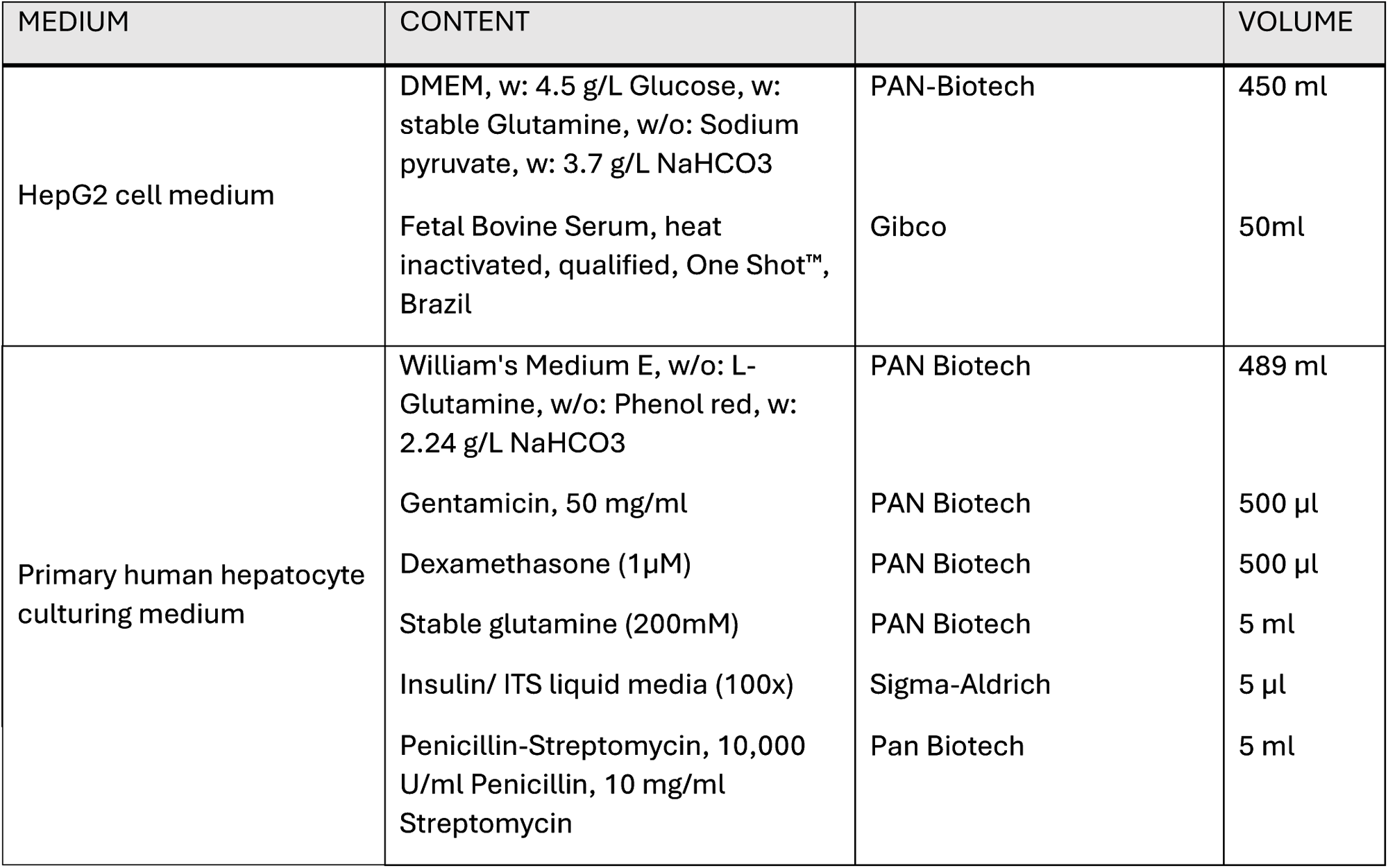
Media

| MEDIUM | CONTENT |  | VOLUME |
| --- | --- | --- | --- |
| HepG2 cell medium | DMEM, w: 4.5 g/L Glucose, w: stable Glutamine, w/o: Sodium pyruvate, w: 3.7 g/L NaHCO3 | PAN-Biotech | 450 ml |
|  | Fetal Bovine Serum, heat inactivated, qualified, One Shot™, Brazil | Gibco | 50ml |
| Primary human hepatocyte culturing medium | William's Medium E, w/o: L- Glutamine, w/o: Phenol red, w: 2.24 g/L NaHCO3 | PAN Biotech | 489 ml |
|  | Gentamicin, 50 mg/ml | PAN Biotech | 500 µl |
|  | Dexamethasone (1µM) | PAN Biotech | 500 µl |
|  | Stable glutamine (200mM) | PAN Biotech | 5 ml |
|  | Insulin/ ITS liquid media (100x) | Sigma-Aldrich | 5 µl |
|  | Penicillin-Streptomycin, 10,000 U/ml Penicillin, 10 mg/ml Streptomycin | Pan Biotech | 5 ml |

**Table S5:** Antibody concentrations

| ANTIBODY | DILUTION |
| --- | --- |
| Mouse $\alpha$ PanRas | 1:1000 |
| Mouse $\alpha$ RAB18 | 1:250 |
| Mouse $\alpha$ GFP | 1:1000 |
| Rabbit $\alpha$ GAPDH | 1:20,000 |
| Rabbit $\alpha$ LC3B | 1:500 |
| Rabbit $\alpha$ RAB18 | 1:500 |
| Rabbit $\alpha$ GFP | 1:7500 |

## Supplementary Figure Legends

**Figure S1.**
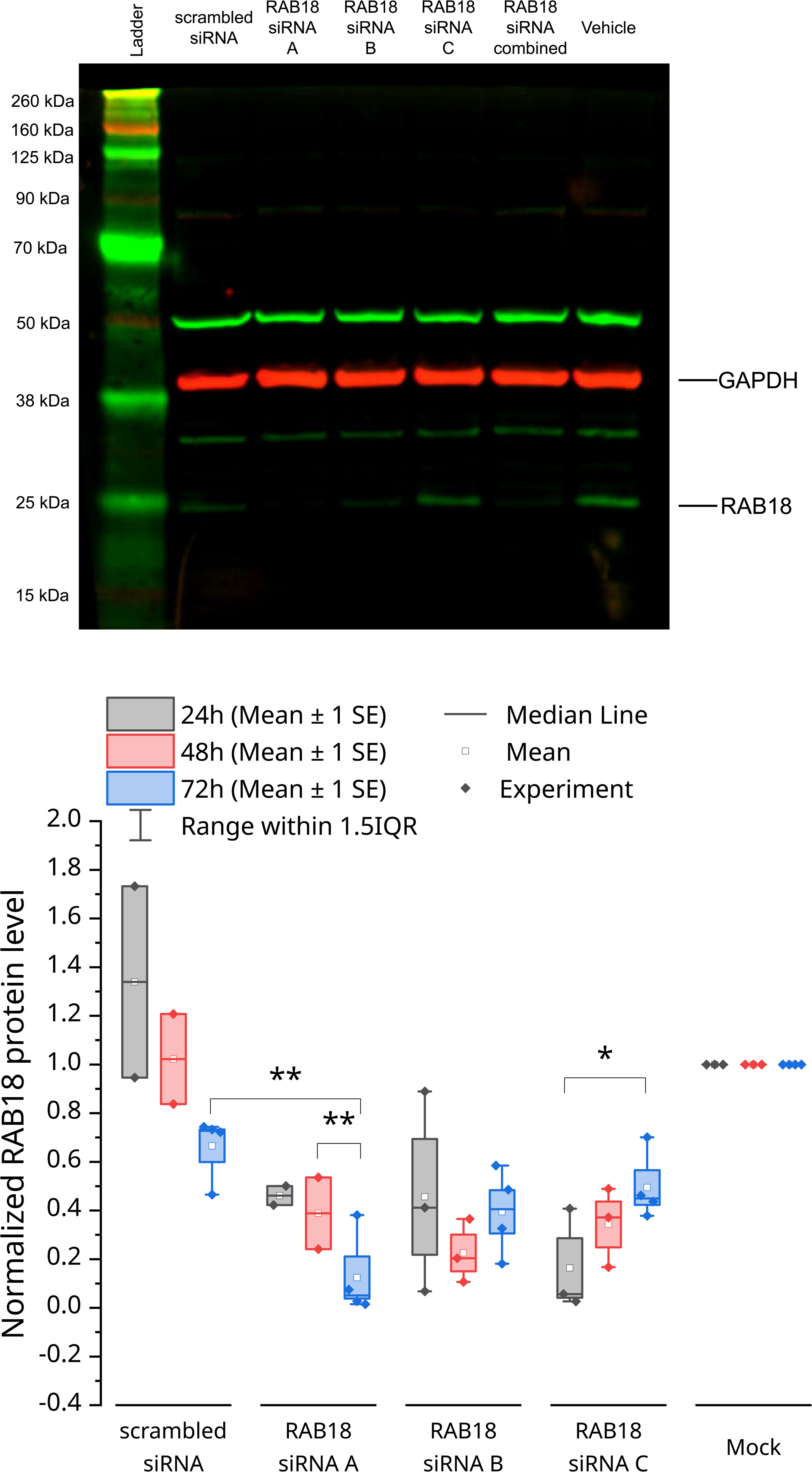
Verification of RAB18 downregulation in HepG2 cells. Representative dual color western blot detecting GAPDH and RAB18 levels in Hepg2 cells 72h post transfection with three separate siRNA or all combined. Quantification of mean intensity normalized to vehicle transfection revealed 88%% knockdown after 72h. N_24h,48h_=3; N_72h_=4. Two 253 sample T-test *p<0.05, **p<0.01.

**Figure S2.**
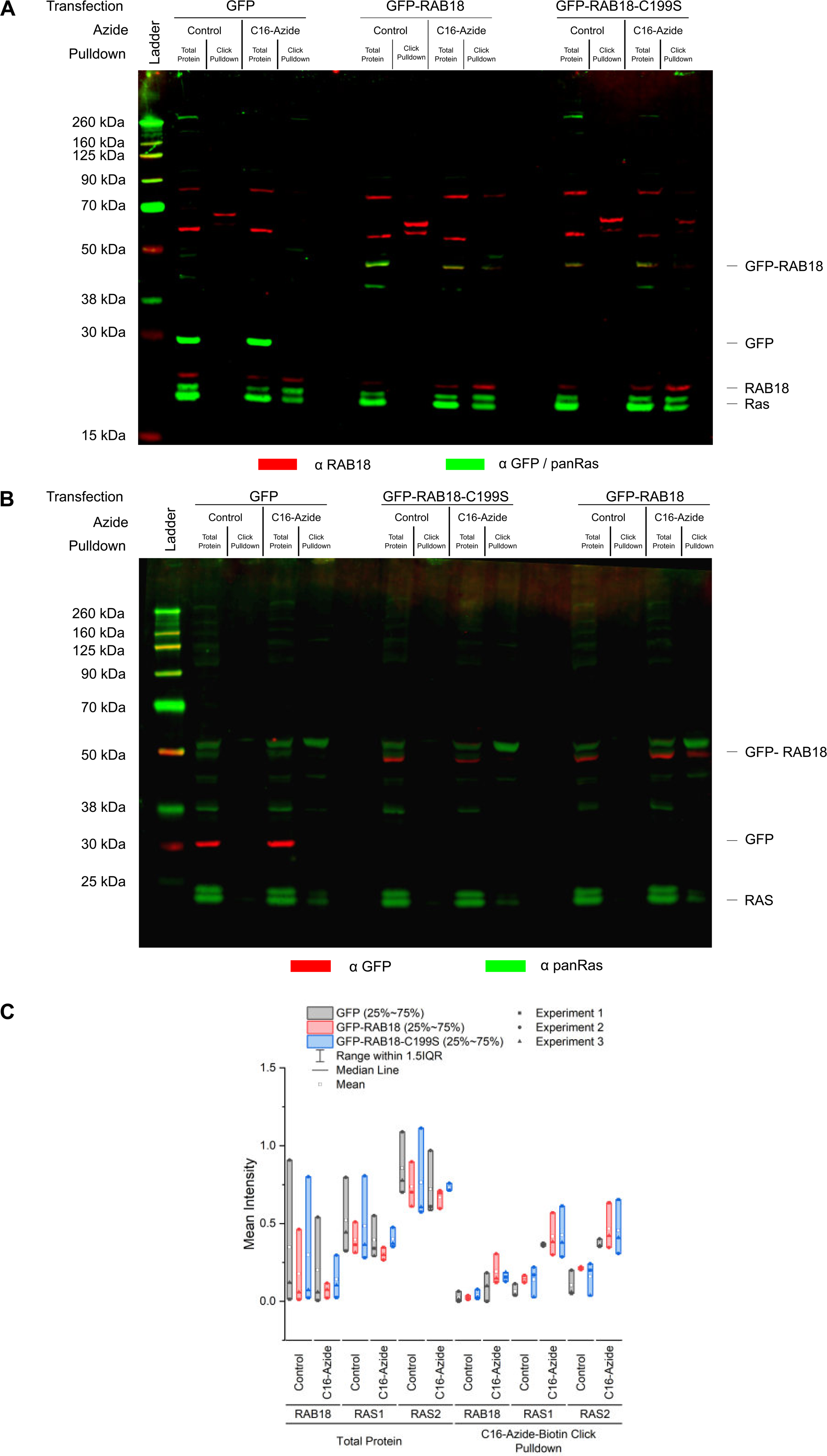
Display of full western blots after azid-linked palmitoyl pull down. **A.** Full dual color western blot of the blot depicted in Figure 3A. Protein amount in lanes (µg): Total Protein: 10, 10, 10, 10, 10, 10. Protein amount in pull-down lanes is the completely recovery from 300, 300,65 (Control GFP-RAB18 transfected), 300,300, 300 input protein (µg) respectively. **B.** Dual color western blot only stained for GFP and panRas to detect GFP RAB18. GFP-RAB18 was detected in the palmitoylated protein pulldown fraction whilst GFP RAB18-C199S was not. **C.** Integrated density of each lane in the three blots used to calculate fold enrichment by click-chemistry pulldown depicted in Figure 3A arranged by transfection. Transfection did not alter the levels of RAB18, RAS1 or RAS2.

**Figure S3.**
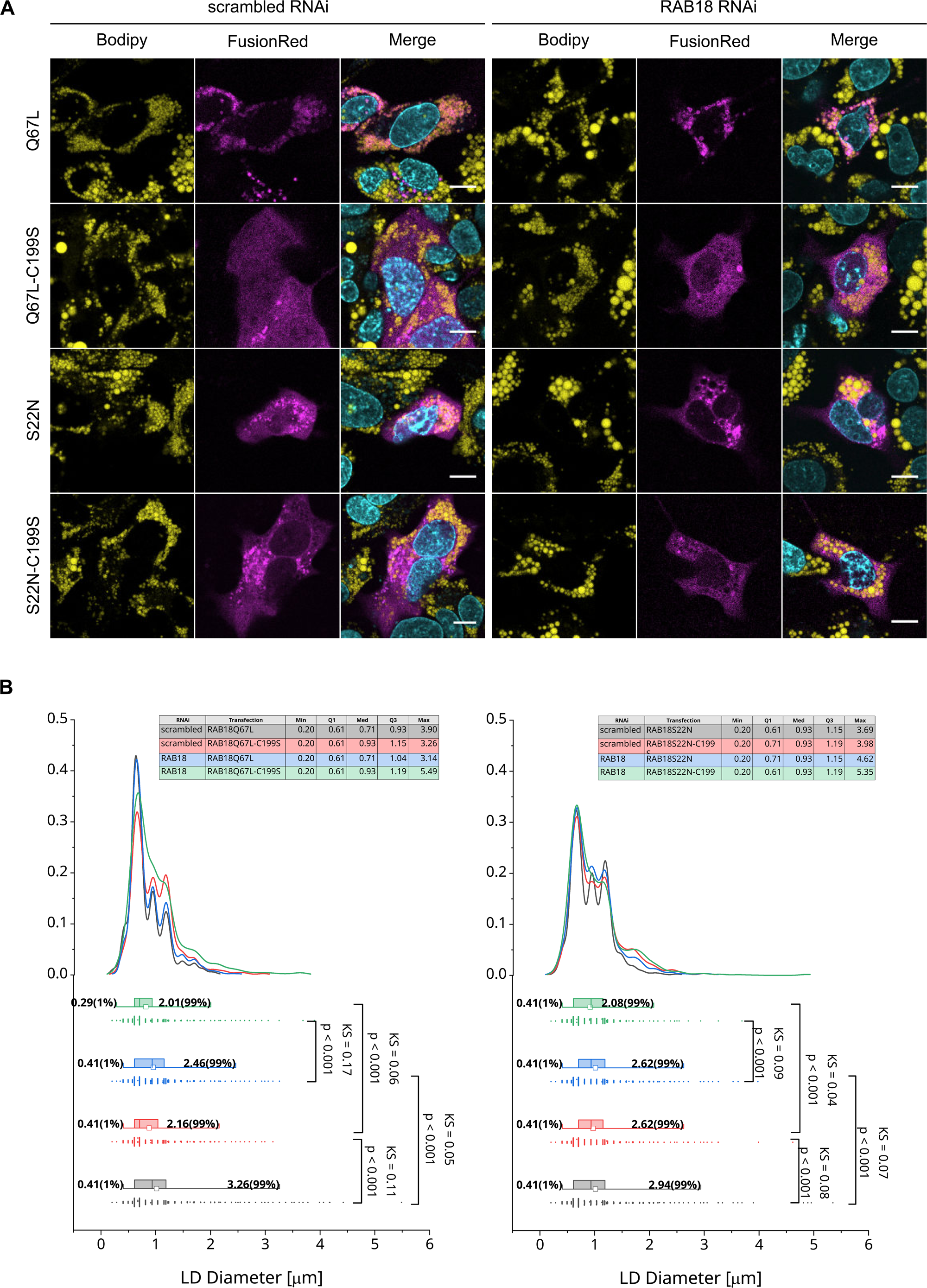
Activity mutants deficient for palmitoylation sites do not localize to LD membrane. **A.** Representative images of HepG2 cells expressing the activity mutants Q67L and S22N alongside their palmitoylation mutants Q67l-C199S and S22N-C199S in cells with RAB18 depletion. **B.** KDE of cells expressing Q67L,Q67L-C199S, S22N and S22N-C199S. Q67L-C199S, S22N and S22N-C199S did not rescue the LD size in cells with RAB18 depletion. Number of experiments (N) and Total number of LDs quantified (n): n, N_Q67L,-scrambled_=4630,3; n, N_Q67L,siRNA_=3914,3; 270 n, N_Q67L-C199S, scrambled_=2662,3; n, N_Q67L-C199S,-siRNA_=1965,3; N_S22N,scrambled_=2393,3; N_S22N,siRNA_=1712,3; N_S22N-C199S-_ 271 _siRNA_=2023,3; N_S22N-C199S,-siRNA_=1709,3

**Figure S4.**
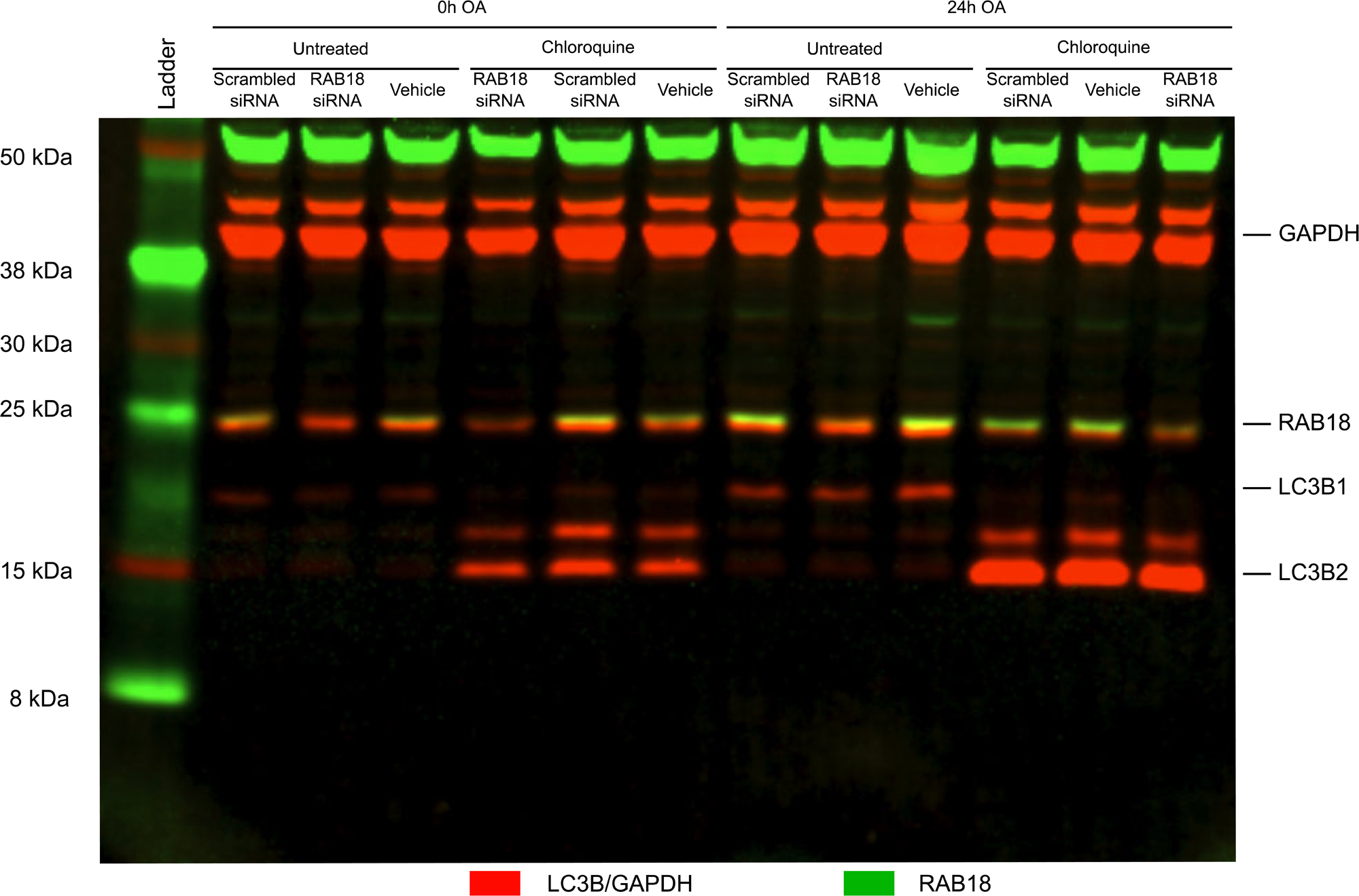
LC3B Westernblot. Full representative dual color western blot of HepG cells with and without RAB18 depletion detecting LC3B, RAB18 and GAPDH displayed in Figure 4C.

**Figure S5.**
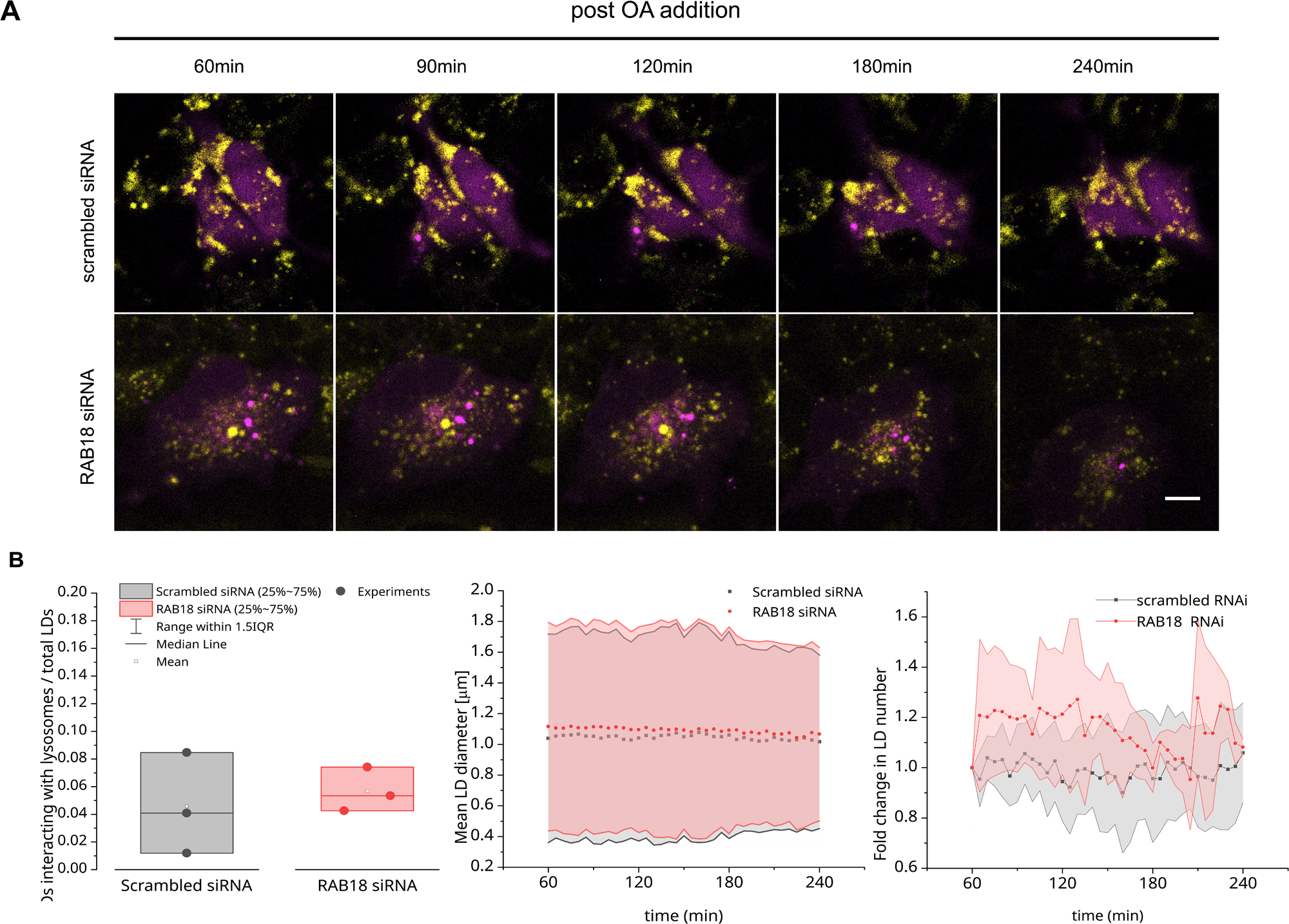
RAB18 depletion does not affect autophagosome-LD interactions after induction of LD formation. **A.** Representative timeseries of HepG2 cells with RAB18 depletion expressing CFP-LC3B (magenta) covering 4 hours of LD formation stained with BODIPY493/503 (yellow) starting 1h post OA addition. **B**. The fraction of LDs interacting with autophagosomes, LD mean area and the formation of new LDs were not altered in cells with RAB18 depletion. N=3. Scalebar: 25µm

**Figure S6.**
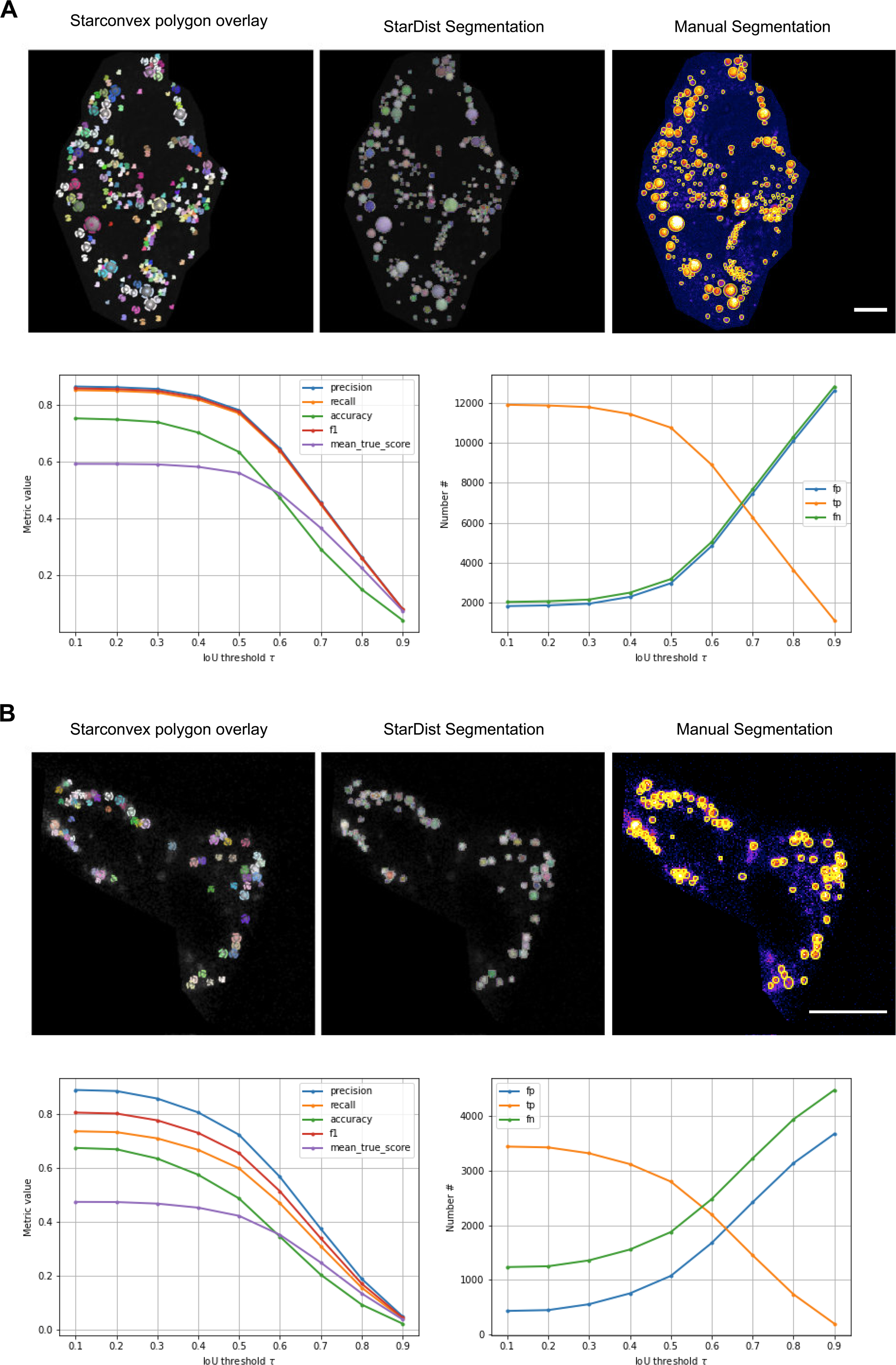
Segmentation with star convex polygon-based machine learning algorithm StarDist. **A.** Exemplary comparison of manual annotated LDs detected with FCARS at 2847cm^-1^ and automatic segmentation obtained by using a custom model trained with the StarDist machine learning algorithm. Evaluation of precision recall, f1 and accuracy revealed a high precision and recall for low Intersection of Union (IoU) thresholding decreasing drastically above an IoU tau of 0.5. **B.** Comparison Analogous to A for LD detected with BODIPY staining. Comparison with annotated data reveals the same pattern as seen with LDs detected with CARS. Overall detection of LDs with BODIPY had a higher false negative (fn) rate than with CARS detection, whilst false positive (fp) remained comparatively low. Number of images (N) and Total number of LDs quantified (n): n,N_CARS_= 14032, 39; n,N_BODIPY_= 3855, 37 Scalebar: 25µm

**Figure S7.**
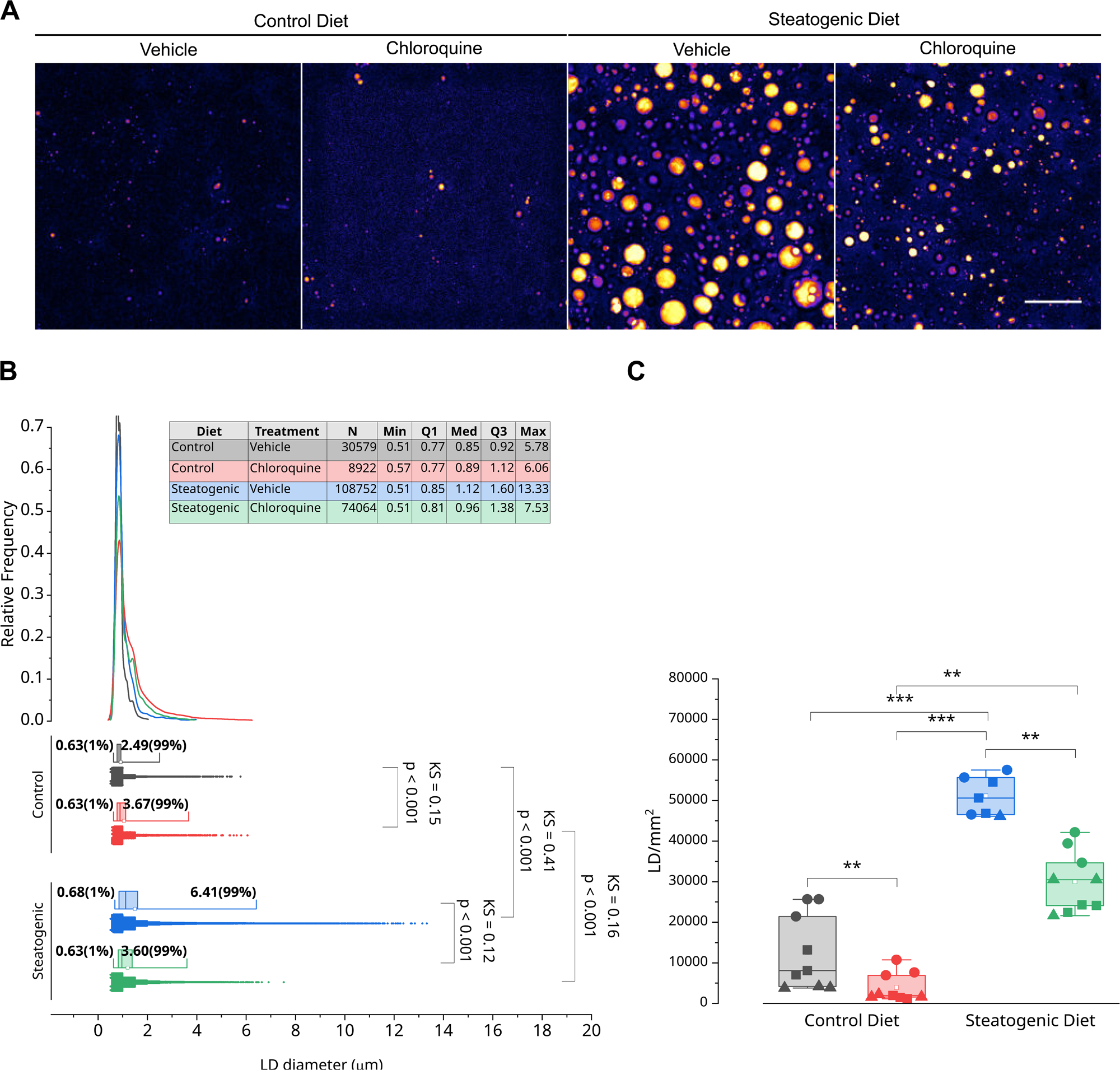
Analysis of cryopreserved slides. **A.** Representative FCARS images of cryopreserved-slides of liver tissue at 2800cm^-1^. Tissue was preserved after 4 weeks of feeding Steatogenic diet and simultaneous Chloroquine injection. **B.** Quantification of the relative LD size distribution of LD detected cryopreserved-liver cuts. Number of mice (N) and Total number of LDs quantified (n): n,N_ControlDiet, Vehicle_ = 30579, 3; n,N_ControlDiet, Chloroquine_= 8922, 3; n,N_SteatogenicDiet, Chloroquine_= 108752, 3; n,N_SteatogenicDiet, Chloroquine_= 74064, 3 **C.**Quantification of LD number per mm^2^ detected in cryopreserved liver cuts. Chloroquine treatment reduced LD number in livers of mice set on a steatogenic or a control diet. Number of mice (N) and Total number images (n): n,N_ControlDiet, Vehicle_= 9,3; n,N _ControlDiet, Chloroquine_= 9,3, n,N_SteatogenicDiet Vehicle_= 8,3; n,N_SteatogenicDiet, Vehicle_= 9,3. Two-sample Kolmogorov-Smirnov-test. *: p<0.05, **:p<0.01, ***:p<0.00. Scalebar: 302 25µm

## Supplementary Tables

**Table S1:**
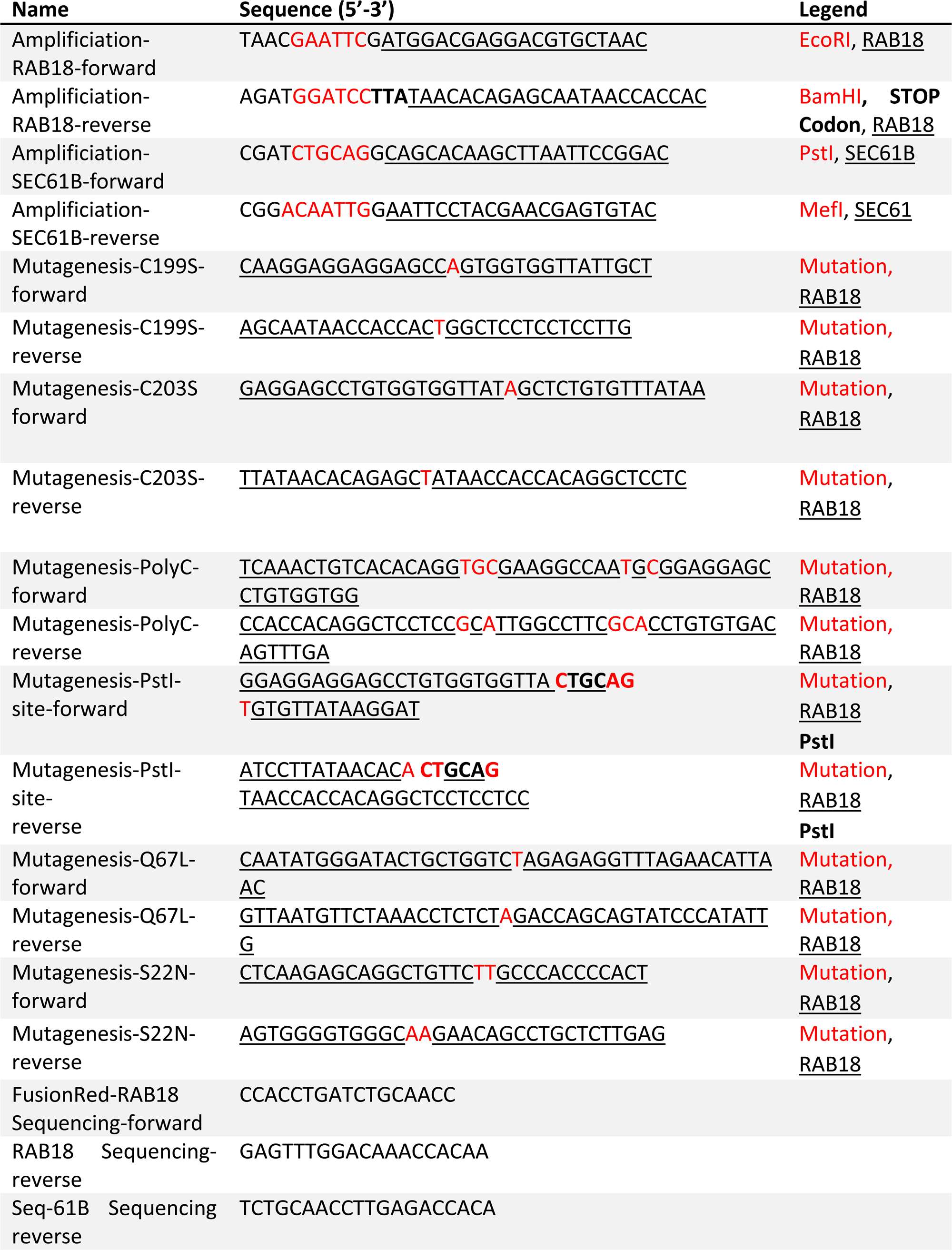
Primer

**Table S2:**
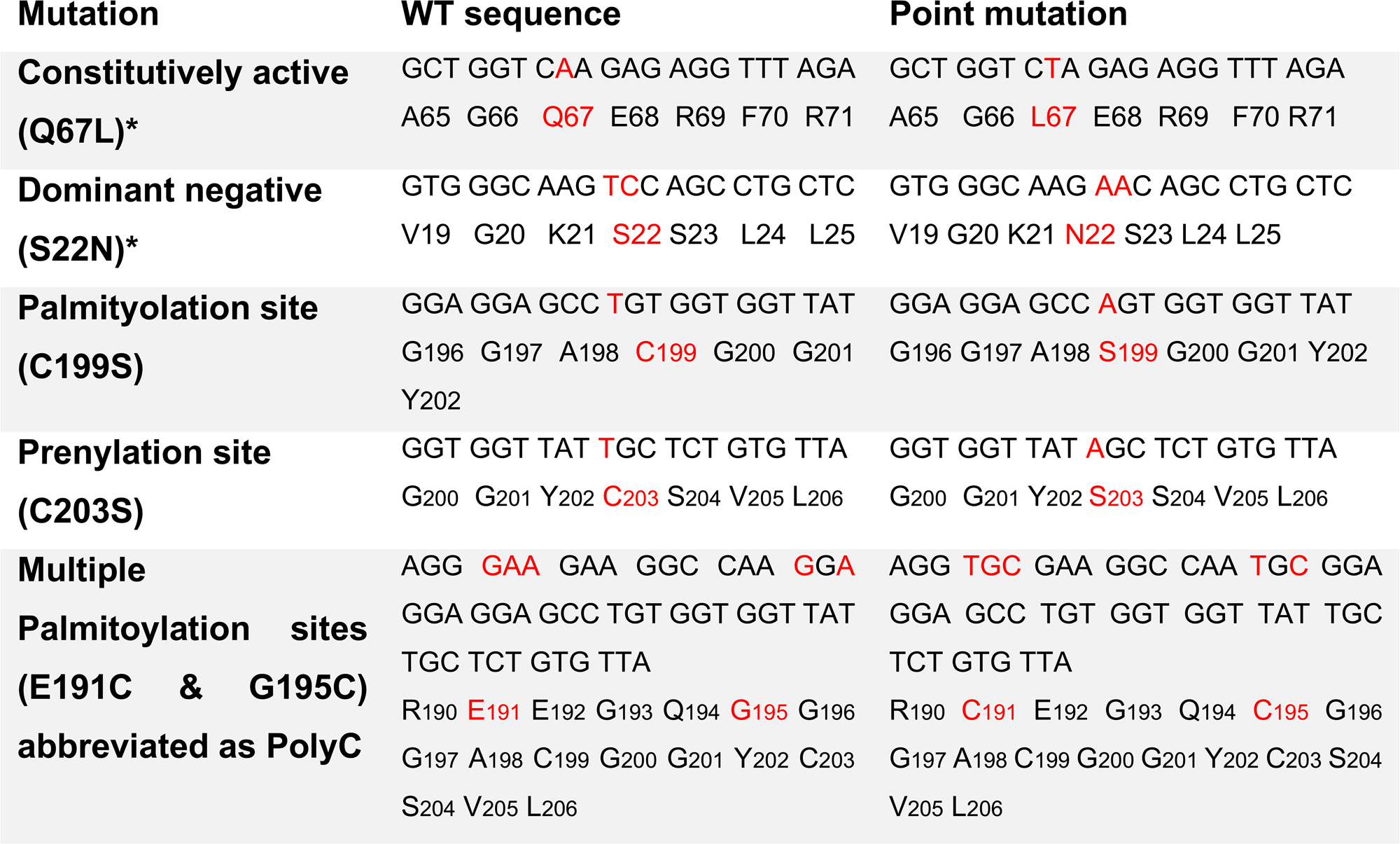
Detailed Mutation overview

## Notes

https://cumulus.ifado.de/published/primetime/Rab18Repository

### Competing Interest Statement

The authors have declared no competing interest.

